# The ontogeny of hippocampal spatial memory is supported by postnatal Cajal-Retzius cells

**DOI:** 10.64898/2026.11.03.763868

**Authors:** I.L. Glærum, R.H.F. Salih, Sachuriga, Y. Roudi, G. Quattrocolo

## Abstract

The functional capabilities of the adult hippocampus depend on specialized circuits that are established during development. Cajal–Retzius (CR) cells persist in the hippocampus long after their disappearance from the neocortex and contribute to the postnatal maturation of hippocampal circuits, but how this developmental role shapes adult hippocampal function remains unknown. Here, we selectively ablated CR cells from the postnatal hippocampus and examined the consequences for spatial learning, navigation, and CA1 spatial representations. CR-cell ablated animals learned reward locations and adapted to changing task conditions, but learned more slowly and made more errors. Reinforcement-learning models suggest that these behavioral differences reflect less stable internal representations guiding behavior. Consistent with this, CA1 spatial representations remained present but were more unstable and showed reduced differentiation of reward locations, particularly when task conditions changed. Our findings identify postnatal CR cells as key developmental contributors to the stability of hippocampal spatial representations and the adaptive behavior they support.

---

The hippocampal region plays a critical role in spatial learning, memory, and flexible navigation [1]. These functions rely on internal representations of space, encoded predominantly in the hippocampus, by place cells [2, 3]. These representations depend on the specialized circuitry of the hippocampus and its interactions with other regions, particularly the entorhinal cortex (EC) [4, 5, 6]. Understanding how this circuitry is established during development, and how its organization gives rise to the functional and computational capabilities of the adult hippocampus, remains a central question in systems neuroscience. A key cell type regulating the development of this circuitry are the Cajal-Retzius (CR) cells.

CR cells are a transient population of glutamatergic neurons that are among the earliest neurons to populate the cortex [7, 8, 9, 10] and are best known for their prenatal role in coordinating cortical lamination through the release of Reelin [11]. Although they rapidly disappear from the neocortex after birth, CR cells persist for months in the hippocampus and medial EC [12, 13, 14, 15, 16], suggesting an additional role in the postnatal development of these regions. Previous manipulations of CR-cell survival have shown impairments in dendritic morphology and memory [17, 18, 19], but these methods lacked the specificity to selectively target CR cells in the postnatal hippocampus. Consequently, it has been difficult to conclusively determine the functional consequences of CR-cell loss. To overcome these limitations, we previously developed a method that combines transgenic targeting [16] and local viral manipulation to ablate CR cells only from the postnatal hippocampus [20, 21]. Indeed, selective postnatal CR-cell ablation alters the maturation of pyramidal-cell dendrites and spines and reduces expression of Latrophilin-2 [20], a molecule involved in establishing EC–CA1 connectivity [22, 23]. Thus, postnatal CR cells contribute to the development of the very circuitry implicated in hippocampal spatial representations. What these developmental alterations mean for the computational capabilities of the adult hippocampus, however, remains unclear.

Using this targeted ablation approach, we sought to understand how the hippocampal circuit develops in the absence of CR cells, and how it differs functionally and computationally from intact hippocampal circuits. Despite the marked cellular and molecular alterations associated with CR cell ablation, CR cell-ablated animals were able to learn the reward locations in an eight-arm radial-arm maze (RAM), and place cells could be observed in CA1. When task conditions were changed by altering the entry location, the animals were able to adapt their behavior. However, clear differences were observed between groups: CR-cell ablated animals learned more slowly and made more errors, both during initial learning and following the change in task condition. To understand the source of these behavioral differences, we fitted reinforcement-learning (RL) agents to the behavior of both groups and compared the properties of those agents. The results suggest that the behavioral phenotype may reflect more unstable internal representations guiding behavior in CR-cell ablated animals. Indeed, electrophysiological recordings from the CA1 revealed a corresponding phenotype: the spatial representations were less stable, and showed reduced differentiation of reward locations, particularly when task conditions change.

Our data demonstrate that postnatal CR cells play a critical role in the development of hippocampal circuitry that can support stable spatial representation while flexibly adapting to changing task demands.

## Results

### Cajal-Retzius cell ablation induces learning deficits in juvenile mice

To determine whether postnatal (P) CR cells are crucial for establishing hallmark functions of the hippo-campus, we took advantage of a double transgenic mouse line that combines Pde1c-cre and NDNF-flox-flp to specifically target CR cells [16]. We performed bilateral injections in the dorsal hippocampi of all animals at birth with a flp-dependent virus (pAAV-mCherry-flp-dta). This injection will only induce apoptosis in cells that express Pde1c-cre^-/+^ and NDNF-flp^-/+^ (CR:DTA+ animals). As control animals do not express cre nor flp (CR:DTA—), CR cells are therefore not ablated, even when infected with the virus. This method effectively ablates 70 to 82% of hippocampal CR cells [20, 21]. Conveniently, the virus induces promoter driven expression of mCherry in all infected cells, allowing us to evaluate the quality of the injection site in both groups. Only animals with successful double-hemisphere injections were included in the analysis (Figure 1A).

**Figure 1:**
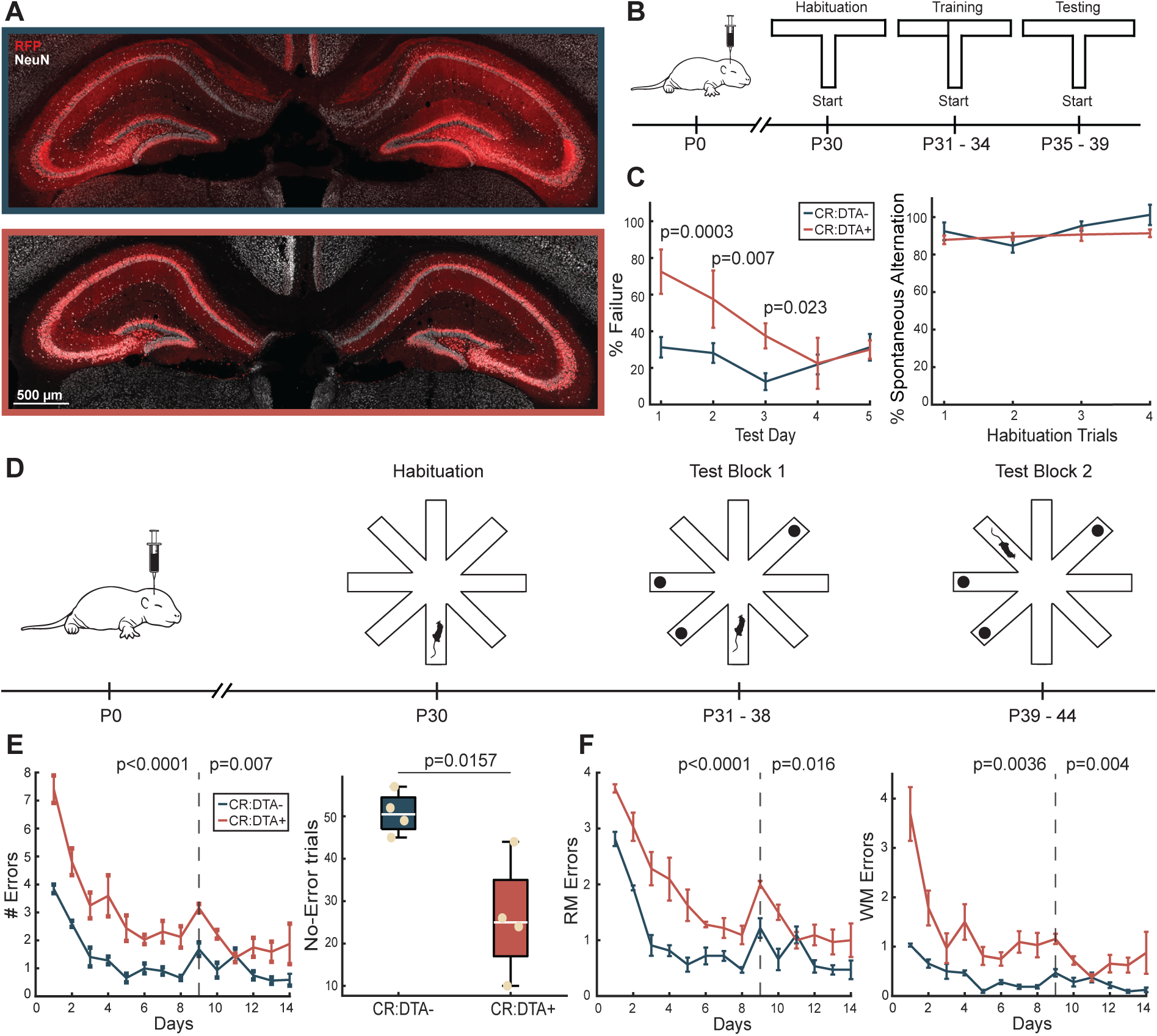
Behavioral experiments highlight learning deficits in CR cell ablated animals. **A**: Representative confocal images of bilateral hippocampal injections. **B**: Timeline schematic for T-maze behavioral testing, bilateral injections at P0, maze habituation at P30 following 4 days of training with forced alteration and 5 days of testing. **C**: Performance of CR:DTA– animals (*blue*), and CR:DTA+ animals (*red*) after training (*left*). Spontaneous alternation scores during habituation trails (*right*). **D**: Schematic of RAM behavioral paradigm, bilateral injections at P0, maze habituation at P30 followed by 8 days in Test Block 1 and 6 days in Test Block 2. Mouse indicating start position. **E**: Average number of errors made per day (*left*). Total number of high-performance trials; no errors made (*right*). **F**: Average number of reference memory errors per day (*left*), and average number of working memory errors (*right*). Dotted line separates Test Block 1 and 2. Bars showing standard error of the mean (SE), and significance assessed by Barnard’s Exact Probability Test, repeated measures ANOVA or two-sample t-test.

We previously demonstrated that postnatal ablation of CR cells reduces dendritic complexity and spine density in CA1 pyramidal neurons [20]. Spine density has been associated with learning and memory processes [24, 25]. High spine density on hippocampal pyramidal cells has been correlated with fewer working memory errors and greater memory capacity of neural networks [26, 27]. As these observations suggest a relationship between spine number and performance in spatial memory tasks, we hypothesized that CR cell ablation may also affect memory.

To test whether CR cell ablation caused memory deficits, we first implemented a baited T-maze test (Figure 1B). We trained P30 mice for 4 days, where the animals were forced to alternate between reward arms. The animals were then tested over 5 consecutive days. During test days, the animal must alternate reward arms without guidance to obtain maximum reward, with a small break between the sample and probe trials (∼2 seconds).

After training, CR:DTA– animals (N=4) already performed the task with 80% accuracy (Figure 1C), while CR:DTA+ animals (N=5) made significantly more mistakes [Π = 0.91, p<0.05]. In fact, CR:DTA+ animals required 3 additional days to reach the same performance level as CR:DTA– animals. This indicates that CR cell ablation caused delayed learning. This impaired performance was not due to deficits in voluntary exploration, as there was no significant difference in their spontaneous alternation during habituation trails [](Figure 1C) nor during exploration of an open field arena [](Sup. 1A-B). The simplicity of the T-maze raises the concern that it primarily tests working memory, as animals only need to remember the arm chosen in the previous trial. To evaluate how CR cell ablation affects both spatial learning and memory beyond simple working memory, we implemented the RAM.

The RAM takes advantage of the innate drive of rodents to locate and retrieve food to assess spatial learning and memory, as it allows evaluation of reference and working memory [28]. Therefore, we tested the spatial memory capabilities of an additional cohort of animals using a variation of the RAM in which three arms were baited.

First, the animals were habituated to the maze at P30 and did not show a preference for any arm (Sup. 1C-F). The animals were then tested the next day in Test Block 1 (TB1). During TB1, the animals were placed in the same starting arm for all trials and tasked with finding the three rewards (Figure 1D). After performing TB1 for 8 days, the animals moved on to Test Block 2 (TB2), where we simply changed the start-arm. This would test whether the animals were able to generalize reward locations across vantage points. The experiment ended after completing 6 days of TB2.

CR:DTA+ animals (N=4) did not reach the same level of performance as CR:DTA– animals (N=4) when navigating the RAM during TB1 (Figure 1E *left*). CR:DTA+ mice made significantly more errors per day than controls [F=27.73, p<0.0001], and they both showed a peak in errors on the first day of TB2. This suggests a reduced ability to generalize from one entry point to another and may indicate that their performance is highly susceptible to external changes to the task. Furthermore, CR:DTA– animals had significantly more trials where they made no errors [t=3.34, p=0.0157] (Figure 1E *right*).

Errors made by both groups consisted of a combination of reference memory (RM) and working memory (WM) errors. RM errors were defined as the first entry into a non-rewarded arm, whereas WM errors reflected re-entry into an arm that had already been visited. While the difference in total errors during TB1 was initially driven by CR:DTA+ animals making significantly more WM errors than controls [F=16.47, p=0.0036](Figure 1F *right*), the slope of the learning curve was driven by CR:DTA+ animals making significantly more RM errors [F=36.5, p<0.0001](Figure 1F *left*). This indicates that CR:DTA+ animals showed deficits in learning the location of the rewards while also keeping track of which arms they had already visited. During the first days of TB2, the difference in RM errors was attenuated without a second increase in WM errors. Thus, although the change in starting location initially impaired navigation, CR:DTA+ animals did not respond to TB2 as an entirely novel task, suggesting that some knowledge of the task acquired during TB1 was retained.

### High speed but low-performance accuracy in CR cell ablated animals

Despite a clear learning deficit, CR:DTA+ animals learn to navigate the maze eventually. Therefore, we speculated that CR:DTA+ may use a different but less effective strategy to navigate. To explore this, we investigated their behavior during individual trials. For a given animal, the trajectory of arm entries was visualized by marking blocks of time where they occupied an arm, with blocks shaded according to arm identity (Figure 2A). These representative examples highlighted how CR:DTA– animals rapidly conform to a stereotyped pattern of arm entries, which could change flexibly according to task condition. CR:DTA+ animals did not display the same conformity, and we observed rapid, seemingly random entries, particularly toward the end of each trial. Interestingly, CR:DTA+ animals exhibited a significant increase in speed [F=104.45, p<0.0001](Figure 2B), which emerged only after task exposure, as no differences were observed during habituation trials (Sup. 2A). This speed difference did not reflect a difference in trial duration during TB1 as CR:DTA+ animals made more errors (Sup. 2B). This increased speed occurred specifically in trials where CR:DTA+ animals made errors [t=-15.55, p<0.0001](Figure 2C) and was absent in high-performance trials [t=0.093, p=0.926](zero errors). While increased speed has been associated with hyperactivity following hippocampal lesions [29, 30, 31], its restriction to goal-directed errors suggests that CR:DTA+ animals may employ alternative navigation strategies rather than exhibiting generalized hyperactivity.

**Figure 2:**
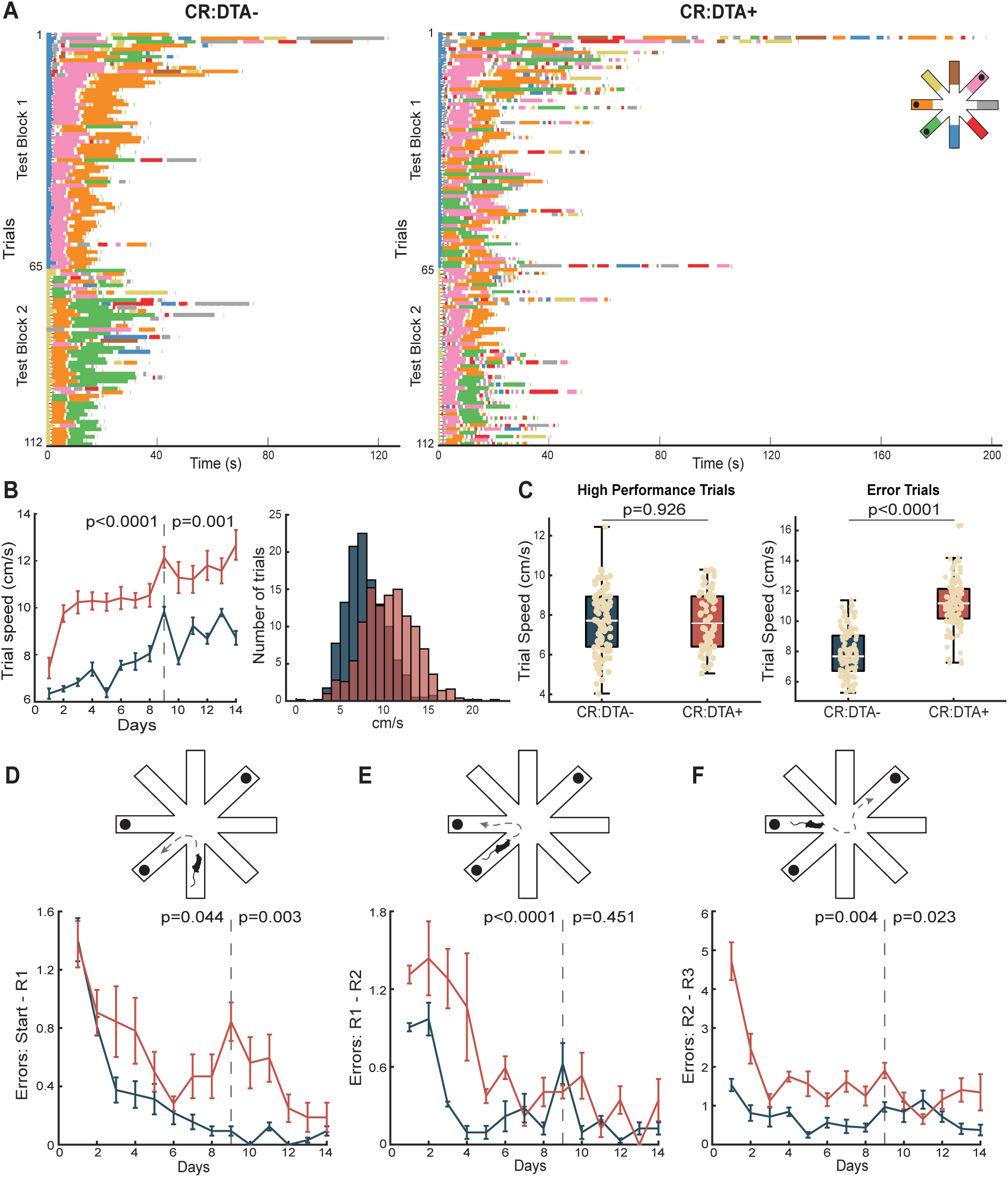
CR cell ablated animals display increased speed but poor accuracy during goal-directed behavior. **A**: Raster plots of arm entries over time for a CR:DTA– animal (*left*), and an CR:DTA+ animal (*right*) colored by arm identity. Note that reward arms are depicted in *green*, *orange* and *pink*. **B**: Trial averaged speed across Trial Block 1 and 2 (*left*), and the distribution of speed for the CR:DTA– (*blue*) and CR:DTA+ (*red*). **C**: Trial averaged speed during high-performance trials (*left*), and error trials (*right*). **D**: Schematic (*top*), and errors made (*bottom*) between trial onset and sampling reward 1 (R1). **E**: Schematic (*top*), and errors made (*bottom*) between sampling reward 1 (R1) and reward 2 (R2). **F**: Schematic (*top*), and errors made (*bottom*) between sampling reward 2 (R2) and the final reward (R3). The order in which rewards were sampled does not matter. Bars showing SE, and significance assessed by repeated measures ANOVA or two-sample t-test.

To validate this, we quantified the number of errors made before retrieving each reward, independently of the order they were sampled. CR:DTA+ animals made significantly more errors than CR:DTA– animals before retrieving each reward during TB1 [F=5.7, p=0.044|F=57.07, p<0.0001|F=15.75, p=0.004](Figure 2DF). However, the magnitude of the errors made was highest when retrieving the last reward (Figure 2F) and was driven primarily by WM errors (Sup. 2E). This pattern suggests a deficit in spatial working memory capacity, which became more pronounced as task demand increased after the first two rewards were retrieved. Interestingly, upon switching to TB2, CR:DTA+ animals made significantly more errors before retrieving the first reward [F=23.07, p=0.023] (Figure 2D), but performance normalized by the second reward [F=0.65, p=0.451](Figure 2E). As these were primarily RM errors (Sup. 2D), this pattern suggests that CR:DTA+ animals relied on negative feedback to reorient, indicating a reduced ability to adapt to changes in the entry point.

### CR cell ablated animals are biased to use a more exploratory navigational strategy

To investigate the strategies animals used to navigate the maze, we modeled behavior during TB1 using reinforcement learning (RL), specifically Q-learning [32]. At each maze location, the agent could choose between available actions, with Q-values representing their expected future reward estimated from previous experience via the Q-learning algorithm. Q-learning is known to converge to the optimal Q-values, such that selecting actions according to these values maximizes the expected cumulative reward. The algorithm is controlled by two parameters: *β* controls how strongly Q-values influence action selection, with low *β* promoting exploration (stochastic choice; *β* → 0) and high *β* favoring the action with the highest Q-value (deterministic choice; *β* → ∞). After each action, Q-values were updated according to the received reward (1 or -0.1), with the second parameter, the learning rate *ɛ*, modulating the strength of the value-update. By fitting *β* and *ɛ* to each animal’s behavior, we assessed whether the difference in performance reflected a different exploration-exploitation balance or a change in the rate of learning.

For each animal, *ɛ* and *β* were fitted by minimizing the mean absolute deviation between the learning curves of the simulated agents and animals across TB1 (Equation 4). The resulting agents closely reproduced the behavior of both CR:DTA– (Figure 3A) and CR:DTA+ animals (Figure 3B). Optimal fits revealed significantly lower *β* values in CR:DTA+ animals (Figure 3C), indicating more stochastic choices compared to controls. However, we did not find a significant difference between the optimal *ɛ*values (Figure 3D).

**Figure 3:**
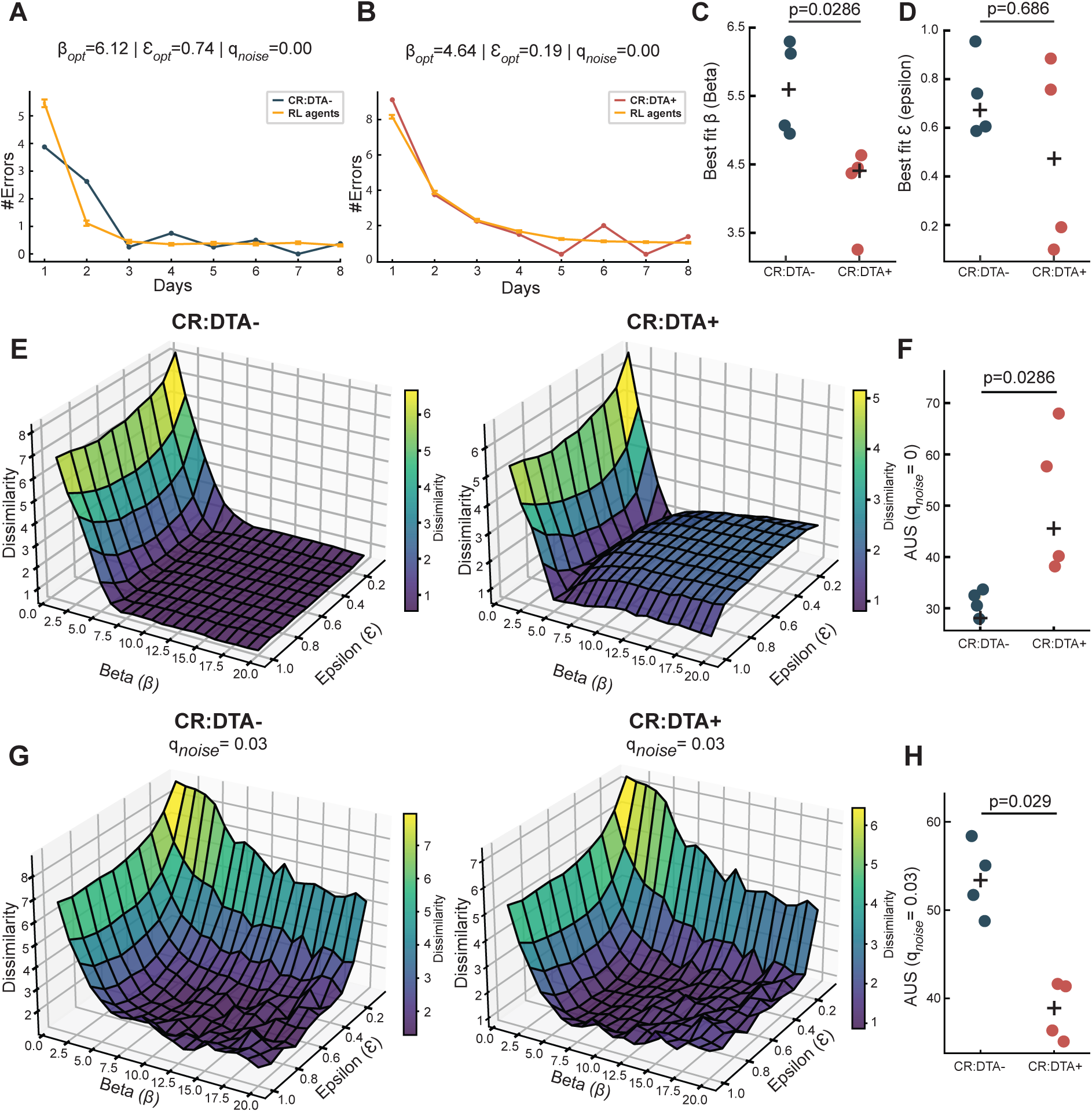
CR-cell ablated animals make more stochastic choices compared to controls. **A**: Example optimized fits of RL agents to the behavior of a CR:DTA– animal in TB1. **B**: Example optimized fits of RL agents to the behavior of a CR:DTA+ animal in TB1. **C**: Box plots of optimal *β* per animal, separated by genotype. **D**: Box plots of optimal *ɛ* per animal, separated by genotype. **E**: Optimization dissimilarity landscape for an example CR:DTA– animal (*left*) and an example CR:DTA+ animal (*right*). **F**: Area under the surface for all animals. **G**: Optimization dissimilarity landscape for the same animals as above using a noisy variant of the RL model with *q*_noise_=0.03. **H**: Area under the surface for all animals using the noisy RL model. In all cases, plus symbol marks the average and significance were assessed using two-sided Mann–Whitney U tests.

To better understand the performance of the RL agents with respect to *β* and *ɛ*, we also looked at the dissimilarity between the learning curves across different values of *β* and *ɛ* (Figure 3E). For the same example animals, low dissimilarity spanned a broad region in CR:DTA– mice but was confined to a narrower region in CR:DTA+ mice (Figure 3F). In parallel to marginal likelihood used in statistical model selection [33], we can employ the area under the dissimilarity surfaces (AUS) to assess model quality without being restricted to the best fit parameters. We found a significantly higher AUS in CR:DTA+ animals (Figure 3F), indicating that their behavior was less robustly captured by the standard RL model.

One assumption of the RL model is that learned action values are represented reliably during learning and updated according to the Q-learning rule. Learning in biological systems is, however, noisy. We thus wondered if the reduced performance of the RL in CR:DTA+ animals can be explained by a noisy learning mechanism. We therefore introduced a noisy RL variant, in which Gaussian noise (*q*_noise_) perturbed learned Q-values, reducing the reliability with which previous experience guided subsequent actions. While a small amount of noise substantially reduced AUS for CR:DTA+ animals, it did not do so for the CR:DTA– animals. In fact, Sup. 3 shows that there is an optimal noise value of *q_noise_* = 0.03 in which AUS is smaller– and thus the noisy RL agent is a better model– for the CR:DTA+ than for CR:DTA– animals; see Figure 3G,H, and Sup. 3). At higher noise levels, however, AUS increased in both groups, indicating that only moderate noise improved CR:DTA+ fits. Thus, robustly capturing CR:DTA+ behavior required a noisier representation of learned action values.

Together, these results show that the strategy of CR cell ablated animals was better explained by an RL model with an impaired ability to retain, and therefore utilize, past experiences. This suggests that CR cell ablated animals are more prone to use a random search strategy to navigate, primarily arising from an inability to maintain a stable representation of the past.

### Postnatal CR-cells are critical to establish a robust spatial representation in the CA1

Postnatal CR-cell ablation led to structural changes in hippocampal pyramidal neurons [20] and reduced spatial information during open field exploration [21]. Here we show that, behaviorally, these animals have navigational deficits consistent with a more stochastic and less value-based strategy. These observations raise the question whether this behavioral instability is reflected in the neural coding of space within the hippocampus.

To test this, we implanted 6-shank high-density silicone probes in the pyramidal cell layer of injected animals that were novel to the task (single units: N_CR:DTA–_=608 | N_CR:DTA+_=593) and recorded while the animals performed the RAM (Figure 4A,B). The RAM protocol remained unchanged except for trial number. To ensure sufficient occupancy coverage of the maze, we increased the number of trials per day (25 trials) and reduced each test block to 3 consecutive days. Implanted CR:DTA+ animals (N=4) showed significant deficits in task performance compared to controls (N=4)[F=9, p=0.008], similar to those of non-implanted animals (Figure 3C). As these were older animals (∼4 months), it confirmed that the deficit in spatial navigation was sustained into adulthood.

**Figure 4:**
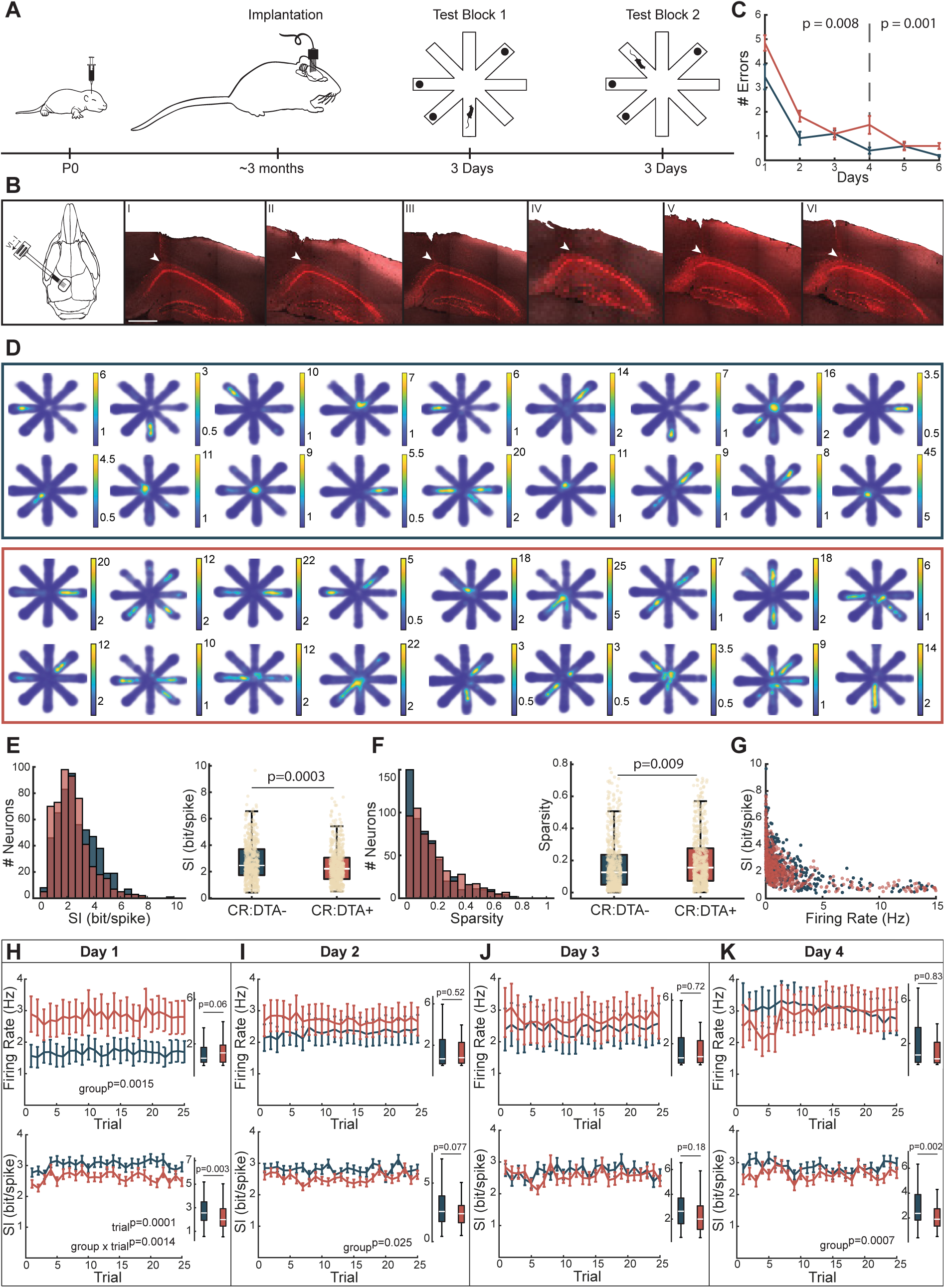
*In Vivo* recordings from implanted mice while freely navigating the RAM. **A**: Schematic of the behavioral set-up with implanted animals. Both TB1 and TB2 have 3 days with 25 trials per day. **B**: Group averaged number of errors per day of the RAM for CR:DTA– (*blue*) and CR:DTA+ (*red*) animals. Tested using repeated measures ANOVA. **C**: Schematic of the implantation site, and traces from shank I-VI of an example animal. **D**: Rate maps from example cells on day 1 in CR:DTA– animals (*top*) and CR:DTA+ (*bottom*) animals. **E**: Histogram (*left*) and box plot (*left*) of the spatial information (SI) for all cells in both groups. **F**: Histogram (*left*) and box plot (*left*) of the sparsity for all cells in both groups. **G**: Depiction of the relationship between firing rate (FR) and SI for all cells. **H**: FR (*top*) and SI (*bottom*) for all trials in day 1. **I**: FR (*top*) and SI (*bottom*) for all trials in day 2. **J**: FR (*top*) and SI (*bottom*) for all trials in day 3. **K**: FR (*top*) and SI (*bottom*) for all trials in day 4. Scale bar: 500 *µ*m. Bars showing SE, and statistics: two-sample t-test or LMM (SI | FR ∼ group × trial, neuron as random effect).

We recorded high-quality units from 6-shank silicone probes (Sup. 4B,C), generated rate maps for each cell (Figure 3D, Sup. 5) and quantified their spatial information content [34]. Postnatal ablation of CR cells resulted in a significant decrease in spatial information [t=4.17, p=0.0003](Figure 4E) and a significant increase in sparsity [t=-2.58, p=0.009](Figure 4F). These effects on spatial information were not driven by differences in firing rate, as spatial information values were not clustered around neurons with extremely high or low firing rates (Figure 4G). Furthermore, CR:DTA+ animals had significantly lower spatial information in both high-performance and error trials (Sup. 4D,E). This confirms the impairment in spatial representation of CR cell ablated animals previously reported in open field experiments [21].

CR cell ablation has previously been shown to increase firing rates of CA1 pyramidal cells during open field exploration [21]. When we compared the firing rates during RAM, we observed no group differences in the average firing rate per day. However, from trial to trial the firing rates differed significantly between the groups on day 1 [F=23.18, p=0.0015](Figure 4H). Additionally, spatial information was also significantly reduced in CR:DTA+ animals, both when averaged per day and when examined per trial on day 1 [t=2.99,p=0.003], and day 4 [t=3.16, p=0.002](Figure 4H,K). Interestingly, spatial information was also lower on day 2 in CR:DTA+ animals despite comparable firing rates [F=5.02, p=0.025](Figure 4I). This indicates that spatial information deficits are not simply a consequence of the altered firing rate. Instead, the trial-to-trial variability in the firing rate observed on days 1 may reflect a reduced stability of the spatial representations during initial learning (day 1) and transfer learning (day 4).

### CR cell ablation results in an unstable spatial representation

As spatial memory performance has been linked to the stability of place fields [35], we sought to investigate whether the reduced spatial memory performance observed in CR cell-ablated animals (Figure 4C) could be explained by an instability in their spatial representations. To assess the stability of each cell’s spatial firing, we computed trial-to-trial correlations between rate maps. This revealed cells with consistently high correlations, indicating stable spatial firing across trials, in both groups (Figure 5A). However, a substantial proportion of cells in the CR:DTA+ group exhibited highly variable trial-to-trial correlations, reflecting an irregular stability pattern across trials.

**Figure 5:**
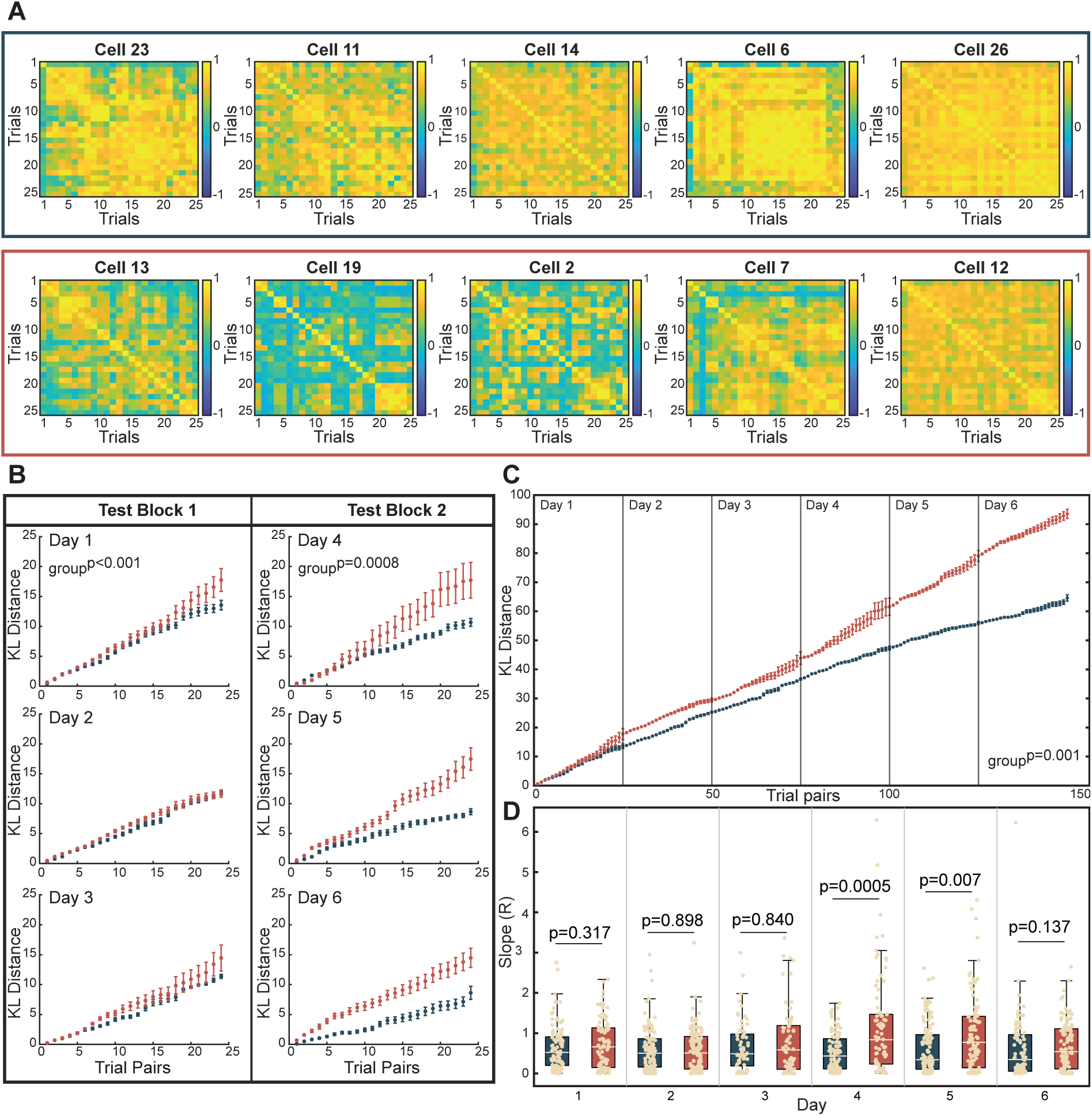
Instabilities in the spatial firing of cells from CR cell ablated animals. **A:** Trial-trial correlations of example cells from CR:DTA– (*top*) and CR:DTA+ (*bottom*) animals. **B**: Group averages of the KL distance calculated on executive trial pairs in each recording session from day 1-3 in TB1 (*left*) and day 4-6 in TB2 (*right*). **C:** Continued cumulative median KL distance from each animal in B. This assumes each animal median captures a population trend. **D:** Quantification of the slope (R: linear regression) of cumulative KL distance of each cell for each day. Bars showing SE, and statistics: LMM (KL divergence ∼ group × session, animal as random effect).

To quantify this observation, we calculated the Kullback-Lieber (KL) cumulative distance for all cells per day between consecutive trial pairs. The KL distance (or relative entropy) is a metric used to quantify the difference in information represented by two distributions with high values indicating greater difference. In this case, the KL distance is calculated for each cell between trial *t* and trial *t*+1 (trial pairs). This gives a single cell measure for the change in spatial firing from one trial to the next. Importantly, we only included spatial bins that had been visited in both trial pairs to avoid any occupancy bias. CR:DTA– animals had significantly smaller cumulative KL distance compared to CR:DTA+ animals on days 1 and 4 (Figure 5B). Over all 6 days, considering each animal per day, the CR:DTA+ animals had significantly higher KL distance compared to CR:DTA– animals [Full model: estimated difference = 1.67 bit, 95% CI [0.68, 2.67], t(1199) = 3.31, p=0.00097] (Figure 5C). To understand the rate at which each cell changed their spatial firing, we looked at the slope of the cumulative KL distance for each cell per day. The cells from CR:DTA+ animals had significantly steeper slopes compared to cells from CR:DTA– animals (Figure 5D), showing a higher rate of change on days 4 and 5.

The absolute KL distance values were higher in cells from CR:DTA+ animals on days 1 and 4, and their rate of change was also higher. This indicates that the spatial representation of CR:DTA+ animals was more unstable, particularly during the initial learning phase (day 1) and after task manipulation (day 4). Both groups showed a reduction in KL distance during days 2 and 3, coinciding with improved task performance, suggesting that the spatial maps stabilized with learning. However, when the entry arm was altered (TB2), the CR:DTA+ group showed a pronounced increase in instability, while cells from the CR:DTA– animals maintained stable representations, indicated by a relatively flat KL distance slope. Cells from CR:DTA+ animals also had significantly higher KL distance values on day 5, suggesting that their spatial maps remained unstable despite improvement in task performance. This highlights a critical change in neural representations when the task transitions from TB1 to TB2, which CR cell ablated animals adapt to less effectively.

### Reward responses are susceptible interference in CR cell ablated animals

The instability of the spatial maps in the CR:DTA+ animals may explain why they make more RM errors (Figure 1F), impairing their ability to rely on a stable structure to navigate. However, we also observed that the CR:DTA+ animals have a deficit in WM, as they make more WM errors when retrieving the final reward (Figure 2F), indicating that they lose track of where they have been. This observation was supported by RL agents capturing CR:DTA+ mouse behavior better with more unstable value representations (Figure 3I). Therefore, we hypothesize that CR:DTA+ animals might not be able to sufficiently differentiate the reward zones to map their specific location in the environment.

To test this, we investigated neural responses before and after entering reward zones 1-3 (Figure 6A). Spikes were binned into 20 millisecond time bins and aligned to each reward zone within an 8-second time window before and after the reward zone entry.

**Figure 6:**
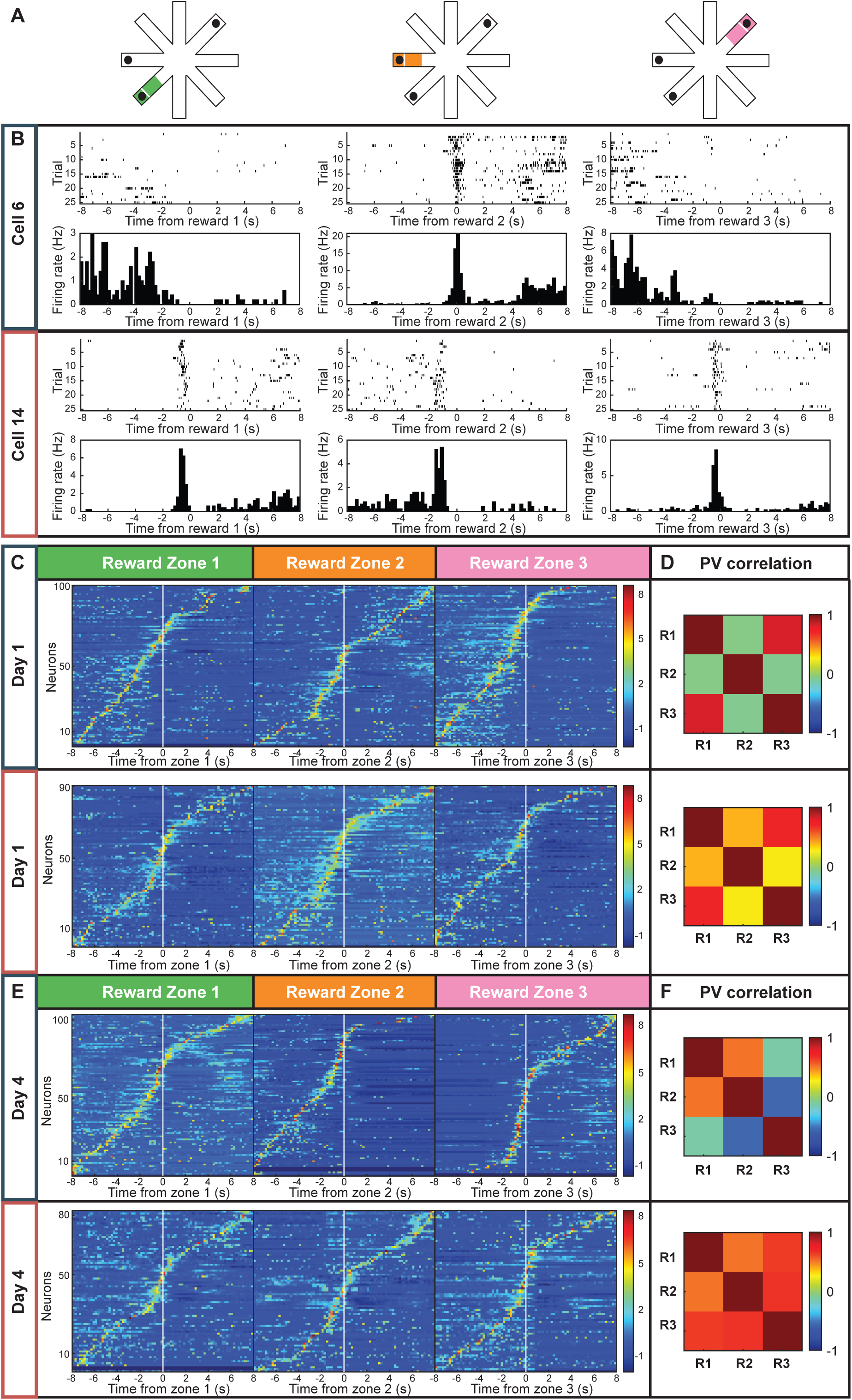
Neural responses to the entering of reward zones. **A**: Schematic of reward zones 1 (*green*), 2 (*orange*) and 3 (*pink*). White lines depicting point of alignment for the following panels. **B**: Raster plots and peri-event histograms of an example cell from a CR:DTA– animal (*top*) and from an CR:DTA+ animal (*bottom*) aligned to reward zone 1 (*left*), 2 (*middle*) and zone 3 (*right*). **C**: Z-scored population responses from a CR:DTA– animals (*top*) and CR:DTA+ animals (*bottom*) on day 1. Neurons are sorted by their peak responses aligned to reward zone 1 (*left*), 2 (*middle*), and zone 3 (*right*). **D**: Population vector (PV) responses correlated over reward zones (R1-R3) for CR:DTA– (*top*) and CR:DTA+ animals (*bottom*) on day 1. **E**: Z-scored population responses from a CR:DTA– animals (*top*) and CR:DTA+ animals (*bottom*) on day 4. Neurons are sorted by their peak responses aligned to reward zone 1 (*left*), 2 (*middle*), and zone 3 (*right*). **F**: PV responses correlated over reward zones (R1-R3) for CR:DTA– (*top*) and CR:DTA+ animals (*bottom*) on day 4.

We observed diverse neural responses when animals entered the three reward zones. Although many cells in both groups responded to multiple zones, CR:DTA– animals exhibited more cells with a single peak, whereas CR:DTA+ animals showed a high proportion of cells with peaks across multiple reward zones (Figure 6B). The population activity revealed that both groups tiled all three reward zones on day 1 (Figure 6C). However, population vector (PV) responses to individual zones were more strongly correlated with each other in CR:DTA+ animals than in controls (Figure 6D), indicating reduced differentiation of firing patterns across reward zones. This divergence became more pronounced by day 4, as CR:DTA– animals exhibited sharper tiling around each reward zone (Figure 6E), whereas CR:DTA+ responses remained largely unchanged relative to day 1. Interestingly, in CR:DTA– animals, the PV for reward zone 3 became anti-correlated with those for zones 1 and 2, while in CR:DTA+ animals all three PVs remained highly correlated (Figure 6F). Additionally, we grouped cells with fields in a reward location (Sup. 6; CR:DTA– = 41.2%, CR:DTA+ = 38.9%) and examined their cumulative KL distance,and found that these reward-field cells were stable across TB1 but became unstable in TB2 in the CR:DTA+ group (Sup. 6). This indicates that reward-field cells in CR:DTA+ animals are more susceptible to changes in task conditions, such as a change of entry point.

These results suggest that CR:DTA– animals progressively refine and differentiate their reward zone representations over time, while CR:DTA+ animals show increasing overlap between zones, consistent with destabilized reward representations following the entry-point manipulation in TB2.

### The spatial instability of cells from CR cell ablated animals is linked to a lower capacity to map space onto reward representations

A population of hippocampal pyramidal cells can maintain a stable spatial code even as individual neurons vary in their firing [36, 37, 38, 39]. We therefore asked whether the single-cell instabilities around reward locations were sufficient to disrupt the population-level representation in CR cell–ablated animals. We hypothesized that reward zones remained represented in both groups, but with degraded information about their spatial identity. To investigate this, we trained two support vector machine (SVM) decoders on population activity: one that classified reward vs. non-reward locations, and a second that further distinguished the reward zone identity.

We applied both decoders to all animals using a bootstrap approach in which 17 cells were sampled with replacement on each iteration (x100). Spike trains were binned (40 ms) and smoothed (*σ* = 2 bins) to match behavioral timescales, and decoder performance was estimated using 5-fold cross-validation. Shuffle controls were generated by training on randomized class labels (see Methods).

When decoding whether the animals were in a reward location (Figure 7A), both groups showed high decoding accuracy (Figure 7B, Sup. 7A), though CR:DTA– animals performed significantly better [F=4.3, p=0.0009|F=2.7, p=0.02]. This difference was primarily driven by a drop in accuracy in CR:DTA+ animals on day 4. In CR:DTA– animals, the decision boundary cleanly separated reward from non-reward activity and improved across days 1–4 (Figure 7C, top). Interestingly, CR:DTA+ animals showed good separation on days 1–2, but the Receiver Operating Characteristic (ROC) curve approached the diagonal on day 4 (Figure 7C, bottom), indicating overlapping distributions. Thus, population activity reliably signaled reward occupancy in both groups, except on day 4 in CR:DTA+ animals.

**Figure 7:**
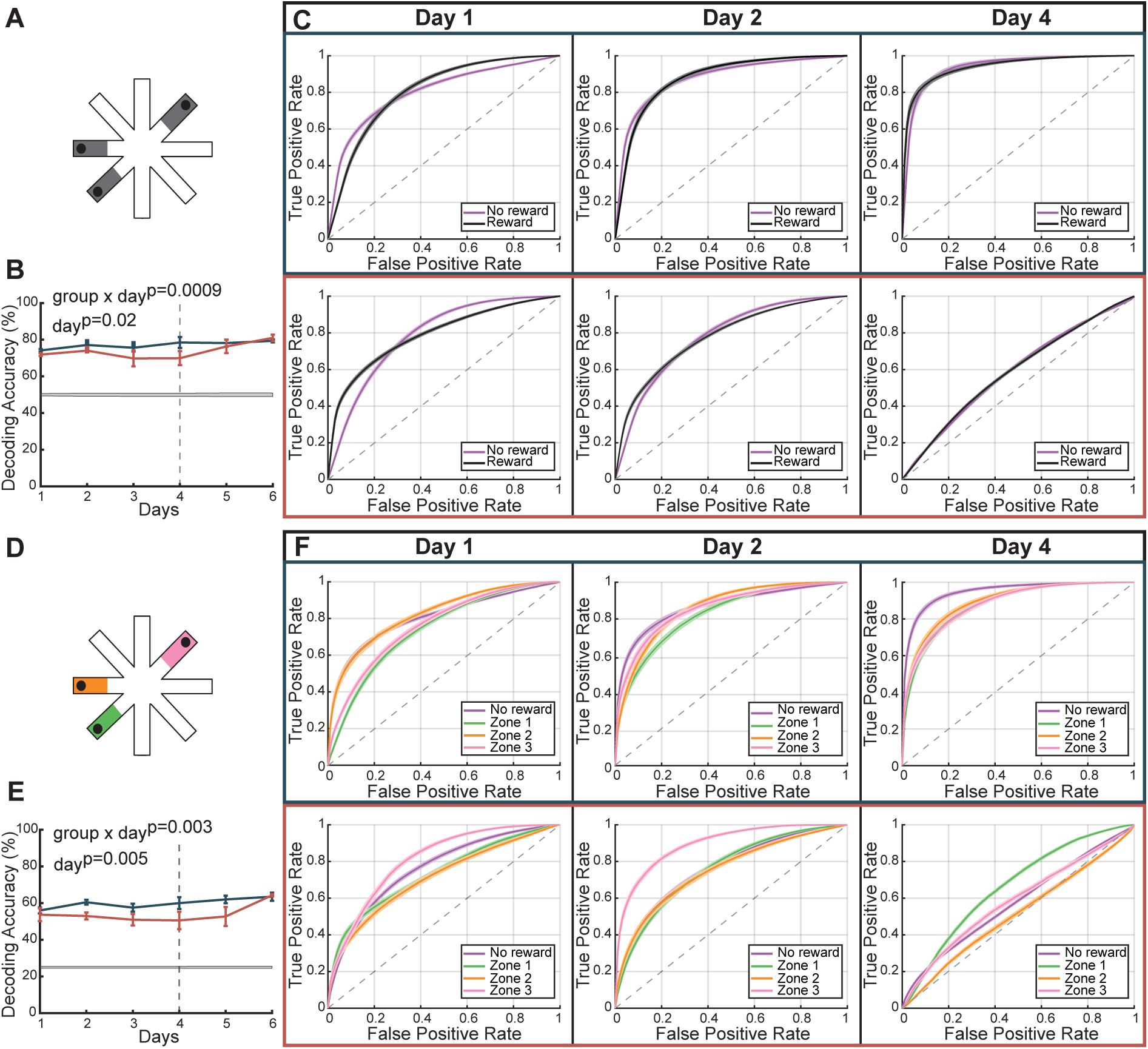
Impaired decoding of reward locations. **A**: Schematic of decoding classes; black representing reward and the remaining being non-reward. **B**: Group averaged decoding accuracies, with corresponding shuffle distributions (*gray*). **C**: ROC curved for decoded classes (reward (*black*) and non-reward (*purple*)) from an example CR:DTA– animal (*top*) and CR:DTA+ animal (*bottom*) from days 1,2, and 4. **D**: Schematic of decoding classes depicting reward zone identities (*green, orange, pink*) and the remaining being non-reward. **E**: Group averaged decoding accuracies, with corresponding shuffle distributions (*gray*). **F**: ROC curved for decoded classes (reward zones 1 (*green*), 2 (*orange*), 3 (*pink*) and non-rewarded (*purple*)) from an example CR:DTA– animal (*top*) and CR:DTA+ animal (*bottom*) from days 1,2, and 4. Diagonal indicated in gray. Bars showing SE, and statistics: LMM (accuracy ∼ group × day, animal as random effect).

Decoding reward-zone identity (Figure 7D) reduced accuracy in both groups, but CR:DTA– animals outperformed CR:DTA+ animals on days 2–5 [F=3.73, p=0.003|F=3.43, p=0.005] (Figure 7E). CR:DTA– animals showed clear and improved class separation (Figure 7F, top), whereas CR:DTA+ boundaries were harder to discriminate (Figure 7F, bottom, Sup. 7B). The decoder appears to generalize a specific reward zone to the other reward classes (Sup. 7), explaining why performance remained high when reward-zone identity was not required.

These results indicate that the population code remains sufficiently able to distinguish reward from non-reward locations, except when CR cell–ablated animals are exposed changing task conditions. However, when increasing the demand on the decoder, the network cannot reliably inform which reward location the animal is occupying. Such reduced reward–zone discrimination could impair the animal’s ability to track which rewards have been obtained within a trial, potentially contributing to the increased errors in retrieving the final reward observed in CR:DTA+ animals (Figure 2F).

## Discussion

The spatial and computational capabilities of the adult hippocampus emerge from circuitry that undergoes substantial maturation during postnatal development. CR cells persist in the hippocampus throughout much of this postnatal maturation, in contrast to their rapid disappearance from the neocortex [16], raising the question of whether their persistence is important for the functional maturation of hippocampal circuitry. Here, we find that while CR-cell ablated animals learn to navigate to rewarded locations, their spatial memory is impaired. CR:DTA+ animals learn more slowly than controls and navigate faster but less accurately under conditions of high task demand. RL models capture this behavioral variability through more stochastic action selection and a reduced ability to reliably maintain learned action values over time. Electrophysiological recordings from CA1 reveal unstable spatial maps, particularly in the representation of reward zones. Together, these findings indicate that postnatal CR-cell ablation compromises the formation of a robust hippocampal spatial code, yielding neural representations insufficient to support effective spatial learning.

Place cells maintain stable spatial representations within familiar environments while flexibly remapping when environmental cues or task demands change [2, 40, 41, 42]. At the population level, spatial coding remains reliable despite changes in individual units [36, 37, 38, 39], and place-field stability is associated with successful task performance [35, 41]. Consistent with this, control animals exhibited more stable representations than CR:DTA+ animals during both initial learning and adaptation to the condition change introduced in TB2. Importantly, this disruption did not preclude learning, as CR:DTA+ animals continued to improve with training. By day 5, the second day of TB2, behavioral performance was comparable between groups despite persistent instability of the underlying spatial representation. Moreover, although both groups developed representations of reward sites, these representations differed in their spatial specificity. While cells from control animals differentiated between individual reward locations, several cells from CR:DTA+ animals displayed reduced precision, encoding for all reward locations. Thus, CR-cell ablation does not abolish spatial learning or reward coding but rather compromises the stability and specificity of the hippocampal representations supporting them.

As CR cells are ablated during a period when spatial behavior is still developing, the behavioral phenotype of CR:DTA+ animals may provide insight into which aspects of mature navigation fail to develop normally. The ontogeny of spatial memory is a prolonged process [43]: recognition of familiar objects appears as early as P14 [44], followed by gross contextual discrimination at P17 [45], whereas precise context discrimination and landmark-based navigation emerge around postnatal week 4 [46, 47]. Adolescent rodents also display a higher exploratory drive [48], suggesting that the maturation of spatial cognition is accompanied by changes in how animals balance exploration with the use of established spatial knowledge. The behavior of CR:DTA+ animals may similarly reflect a less mature navigation strategy. During goal-directed navigation, they ran faster yet made substantially more errors, and RL models indicated a greater reliance on exploration over exploitation. The ability to generalize learned information to novel situations promotes exploitation of existing knowledge over further exploration as seen in adult mice [49]. Notably, CR:DTA+ animals made more errors when required to navigate from a new start location, consistent with reduced generalization of previously acquired spatial information. Their exploratory bias may therefore reflect a reduced ability to exploit an established representation of the environment. Although increased exploration could reflect differences in motivational drive, its convergence with impaired generalization, reduced spatial stability, and weaker differentiation of reward locations suggests that the phenotype is not purely motivational. Instead, CR-cell ablation may impair the formation of stable spatial representations that can be flexibly used to support mature, goal-directed navigation.

We propose that CR cells act as a developmental scaffold that facilitates key maturation processes in the postnatal hippocampus. Although rabies-virus tracing shows that hippocampal afferents, particularly from the EC, target CR cells at P1 [50], glutamatergic input onto CR cells declines drastically over the first postnatal weeks [51]. This transient connectivity, preceding dendritic maturation of CA1 pyramidal cells [52], supports the idea that CR cells may serve as temporary targets for incoming afferents during circuit development. As CA1 place-field stability has been shown to depend on EC input [53], the spatial instability observed in CR:DTA+ animals could, at least in part, reflect impaired integration of EC afferents. Beyond altered EC–CA1 connectivity, the deficit in reward-related spatial coding may also reflect disrupted interactions between the hippocampus and prefrontal circuits involved in goal-directed behavior. In particular, the spatial representations of CA1 neurons projecting to the prefrontal cortex are preferentially stabilized during reward learning [54], while perturbation of hippocampal–prefrontal communication impairs working memory and reduces spatial decoding around reward sites [55]. Thus, the reduced spatial stability and reward-zone decoding of CR:DTA+ animals may reflect broader disruption of circuits coordinating spatial and goal-related information. By transiently supporting the maturation and integration of CA1 pyramidal cells and their afferent inputs, CR cells may therefore contribute to the establishment of hippocampal circuitry whose functional consequences persist long after CR cells themselves have disappeared.

In conclusion, postnatal CR cells are required for the development of hippocampal circuitry capable of supporting stable and flexible spatial representations. Their premature loss does not prevent spatial learning but leaves hippocampal representations less stable and less able to support efficient navigation when task demands change. These findings reveal that the transient postnatal presence of CR cells is important for the functional maturation of hippocampal circuits, with premature loss having lasting consequences for hippocampal computations and spatial behavior.

## Author contributions

I.L.G. and G.Q. conceptualized the study. I.L.G., R.H.F.S., S., Y.R. and G.Q. developed the methodology. I.L.G., R.H.F.S. and S. performed the investigation and curated the data. I.L.G. and R.H.F.S. performed the formal analysis and prepared the visualizations. I.L.G. wrote the original draft. All authors reviewed and edited the manuscript. Y.R. and G.Q. supervised the study. G.Q. acquired funding.

## Online Methods

### Animals

All experiments were conducted in compliance with protocols approved by the Norwegian Food Safety Authorities and European Directive 2010/63/EU (FOTS ID 24847, 28607 and 31016). All mice were housed in enriched environment cages in an inverted 12 h light/dark cycle with food and water *ad libitum* outside the experimental time line. The mice were temporarily food restricted ∼2 hours prior to all behavioral tests, but had free access to water. Pups were separated from the mother at three weeks of age (P21).

Male transgenic heterozygous Pde1c-cre mice (MMRRC B6.FVB(Cg)-Tg(Pde1c-Cre)IT146Gsat/ Mmucd, MMRRC 030708 [56]) were bred with female heterozygous NDNF-flox-flp mice (donated by Dr. Rob Machold, NYU and Dr. Gord Fishell, Harvard University[16, 57]) to generate a double-transgenic mouse line that was previously reported to effectively label CR cells [16]. We chose to treat the double negative genotype (Pde1c-cre^-/-^;NDNF-flp^-/-^) as our control group and named them CR:DTA–, and the double positive (Pde1c-cre^-/+^;NDNF-flp^-/+^) as the experimental group and named them CR:DTA+. The other genetic combinations were not included in any of the experiments. Animals of both sexes were included and balanced. This resulted in a total of 25 animals (females = 13 | males = 12) from 17 litters. All included animals were only exposed to one experimental protocol.

### Viral injection

All pups were subjected to viral injections at P0. These were bilateral injections with a recombinant adeno-associated virus (pAAV-mCherry-flp-dta | modified from Addgene plasmid #58536; http://n2t.net/addgene:58536; RRID: Addgene_58536; [58] by the Viral Vector Core at the Kavli Institure for Systems Neuroscience) expressing Flp-dependent diphtheria toxin A fragment (AAV-DTA). The viral construct is engineered to drive pan-neuronal expression of the fluorescent protein mCherry, enabling easy assessment of injection quality. Furthermore, the virus contains an inverted DTA segment, which is floxed by FRT sites under the control of the EF-1a promoter gene. In the presence of Flp, which is only present in cells that also express Cre, recombination of the FRT sites will result in inversion and activation of the DTA sequence inducing apoptosis. In the absence of Flp, the DTA sequence will remain inactive, but mCherry will still be expressed. Thus, only CR cells will undergo apoptosis in CR:DTA+ animals (carrying both Flp and Cre). We excluded animals that did not have cells that expressed mCherry in the outer blade of the DG in both hemispheres.

Animals were anesthetized with isoflurane (3%) and head fixed in a stereotaxic frame (Koppf) with a custom-made adapter. The skin was stretched using standard lab tape with a diamond cutout. Both hippocampal coordinates were calculated from lambda (AP + 0.8mm | ML ± 1.2mm | Z − 1.22mm) in each individual mouse. A glass pipette (Drummond Scientific Company) was attached to a Nanoject III injector (Drummond Scientific Company) for injections of 48.4 nl (4.5 nl/s) per hemisphere.

### Behavioral paradigm

Two consecutive days prior to the behavioral experiments, the animals were habituated to the room, handling by the experimenter, and the reward (Oatly chocolate milk). To optimize behavioral engagement the animals were food restricted ∼2 hours before experiment onset, but outside that time window the animals could eat *ad libitum*. The experimenter was blinded to the genotypes in all experiments.

### T-maze

Animals were habituated to the T-maze at P30 for 2×5 minutes in the morning and the afternoon (total 20 minutes). The following day we started 4 days of consecutive training with forced alternation. One target arm was blocked while the other lead to a reward arm. After reward consumption the animal was removed and the blocked arm switched to allow sampling of the other reward. The order in which the target arms were blocked was randomized to avoid choice bias. Each day of training was constituted by 8 trial sets, 4 in the morning and 4 in the afternoon (total of 32 trials). At P35, after receiving training, the animals were tested for 5 consecutive days by removing the blocking. If the animal enters the same arm that was previously sampled, it only received one reward and it was considered an error, but if the animal alternated, it would receive both rewards which was considered a correct choice. A total of 8 trial-sets per day, split by a morning session and an afternoon session, yielded a total of 40 test trials per animal. A minimum of two animals were tested at once and alternated to give a break in between trial-sets. Only after the trial-set finished was the maze thoroughly washed with ethanol and made ready for the next animal.

### Radial-Arm Maze

The RAM maze has eight arms projecting radially outward, each of which with a small food bowl at the end that may contain a reward that are not visible from the center. In this RAM variation, three fixed arms contained reward (one drop of chocolate milk), and the animals were tasked with retrieving and consuming all three before the trial ended. Three distinct distal cues with unique shape and size were distributed around the maze, only one rewarded arm had the cue directly behind it. At P30 the animals were first exposed to an open field trial (1×20 minutes) and then habituated to the RAM for 2×10 minutes in the morning and in the afternoon. The following day, animals started Test Block 1 (TB1), where 3 arms were rewarded and the entry arm was fixed (Figure 1D). This was kept constant for 8 trials x 8 days, then Test Block 2 (TB2) starts on day 9 where the rewarded arms remain constant, but the entry point is changed to a new arm. With 8 trials x 6 days for TB2, this gives a total of 112 trials.

Electrophysiological experiments were conducted on naive mice. The structure of the RAM task is largely the same, but to optimize for spatial coverage, there was only one session each day with 25 consecutive trials. The session is one continuous recording, and the animal was resting in a flowerpot in between trials. With more trials per day, fewer days were needed. Therefore, both TB1 and TB2 had 25 trials x 3 days, providing a total of 150 trials per animal.

### Stereotaxic implant surgery

Naive injected mice, 4 months old, underwent stereotaxic surgery to implant Cambridge Neurotech Silicone probes (6-shank ASSY-236 probes). Each shank is spaced 200 *µ*m apart horizontally, containing 10-11 recording sites that are spaced 15 *µ*m vertically and 16.5 *µ*m horizontally. In total, this yields 64 evenly spaced electrodes.

All surgeries were performed at 1% isoflurane mixed with 0.3% oxygen. Following anesthesia, the animals received subcutaneous injections of anti-inflammatory analgesics (Buprenophine 0.03 mg/kg, Meloxicam 1 mg/kg) and saline immediately after induction. After loss of the toe pinch reflex, the animals were head fixed, shaved, and the skin removed after local sedation with Bupivacaine (1 mg/kg). The animals were resting on a heating pad at 37*^◦^*C for the entirety of the procedure. The skull surface was cleaned for connective tissue before implant coordinates were determined by referencing lambda and bregma. After leveling lambda and bregma (DV < 0.1 mm | 4 mm < AP < 4.2 mm), a circular craniotomy was performed around the hippocampal coordinates calculated with referencing bregma (shank 1: AP − 1.8mm; ML + 0.8mm | shank 6: AP − 2.3mm; ML + 1.9mm). An additional craniotomy was performed over the cerebellum for the ground. The craniotomies were then sealed using Kwik-seal, before base plating using optibond mixed with venus curated with UV light. The seal was then removed, and the probe was implanted -0.9 DV from the dura. The probe was manually pre-assembled to a nano-drive and a Mini-Amp-64 head stage (Cambridge Neurotech). All movable parts of the drive were covered with Vaseline (including the probe), before the nano-drive was anchored to the skull using dental cement. The ground wire was attached to the reference screw using conductive silver paint before the head-stage was cemented in place. The total weight after implantation was less than 3 grams.

Immediately after surgery, the mice were moved to a heated chamber (26 *^◦^*C) for 1-2 days depending on the recovery time of the animal. The mice received additional does of Meloxicam every 24 hours and buprenophine every 6-8 hours during recovery. Once the animals regained their pre-surgery weight, pain management was no longer needed (approximately 3 days). The animals were left to recover for a week before we began recording.

### Recording of hippocampal activity

After recovery, the probe was slowly moved into the pyramidal cell layer of the CA1, identified by theta in the local field potential, ripples and a sharp increase of spiking activity. The probe was kept in the position that yielded the maximal number of units throughout the experiment.

The behavior of the animals was captured by a Basler ace2 camera (Basler AG) equipped with far-red filters. The recording platform was illuminated by two Light Bar-45×100-Red (Edmund Optics). The camera captures 50 frames per second and with each frame a TTL signal is sent to the OpenEphys acquisition board for synchronization of the signal streams.

Neural activity was filtered, amplified, multiplexed, and digitized by the Mini-Amp 64 (Cambridge NeuroTech) and recorded by an Open Ephys acquisition board at a sampling rate of 30kHz (0.1-7,500 Hz, x192; 16 bits).

### Spike sorting and curation

Binary recording files were loaded and band-pass filtered between 300 Hz and 6000 Hz. Spikes were then sorted using the spikeinterface.sorters module with the Ironclust algorithm on shanks with up to 11 sites per shank. Automated spike sorting was followed by manual curation in SpikeInterface Phy2. Manual curation on the sorted clusters was performed by a single experimenter by assessing ISI violations, the quality of the waveform and separation in projected three-dimensional principal component space. Occasionally units were split if judged to contain different waveforms. Data from each session, a set of 25 trials and rest periods, were spike sorted together. Only well isolated units (median waveform amplitude > 40 *µ*V | ISI violation ratios < 0.2 | 30 Hz > average firing rate > 0,05 Hz) were retained for future analysis.

Based on our previous observations of physiological alterations in pyramidal cells following postnatal CR cell ablation [21], we included all cells with an average firing rate below 15 Hz during the task. This threshold excludes putative interneurons with high firing rates in they pyramidal cell layer while retaining other cell types, allowing us to examine how the CA1 network responds to task learning.

### Postprocessing of electrophysiological data

Videos were analyzed using DeepLabCut [59] with a custom trained model established in the lab [21]. The model was trained to identify twelve keypoints (snout, left ear, right ear, neck, back 1, back 2, back 3, back 4, tail 1, tail 2, tail 3, and tail 4) in approximately 3,000 frames. The coordinates of the midbrain were used as the final position coordinates.

For each individual recording session only units that passed the quality metrics were included. Raw position data and spikes times were aligned with the trial start/stop times derived from TTL-synchronized behavioral logs.

Each arm was reconstructed from the Fiji delineated coordinates, converted to arena coordinates, and applied to the position traces. Arm occupancy vectors were generated by assigning position samples to arm boundaries, and short-duration entries/exits (<0.5 s) were smoothed to remove noise (generated from the animal spending time on a zone boundary). Trial-wise position and arm traces were extracted for each session, along with corresponding trial-specific spike times for each cell.

Velocity was computed from the Euclidean distance between successive position samples divided by elapsed time, then interpolated back to the original time resolution. For each trial, trial-specific velocity traces, spike times, and position/arms were stored.

### Perfusion and tissue preparation

For euthanasia, all animals were first anesthetized with isoflurane before they were euthanized with a lethal intraperitoneal injection of pentobarbital (100 mg/kg). After that, the animals were transcardially perfused, first with phosphate-buffered saline (PBS), then with a 4% paraformaldehyde (PFA) solution in PBS (14 ml at a rate of 3.7 ml/min). Brains were harvested and post-fixated for 3 hours in PFA. They were then moved to a 15% sucrose solution overnight (stored at 4°C) and then to a 30% sucrose solution for an additional overnight incubation. Fixated brains were coronally sectioned on either a freezing microtome (ThermoScientific) or a Cryostat (ThermoScientific) in the case of implanted brains.

### Immunohistochemistry

For immunohistochemistry, sections were rinsed in PBS for 10 min and subsequently blocked for 30 min in a blocking solution containing 10% normal donkey serum (Jackson ImmunoResearch) and 0.1% Triton X-100 (Merk) dissolved in PBS. Sections were then incubated with primary antibodies diluted in an incubating solution containing 1% normal donkey serum and 0.1% Triton X-100 dissolved in PBS, overnight for three nights at 4°C with gentle shaking. Primary antibodies were as follows: rat anti-RFP to amplify the mCherry signal, dilution 1:1000 (Proteintech, RF5R); rabbit anti-p73, dilution 1:500 (Abcam, ab40658); guinea pig anti NeuN, dilution 1:1000 (Merk, ABN90P). The secondary antibody staining (directly conjugated Alexa fluorophores, 1:500 diluted in incubating solution) followed after 3×1 h rinsing with PBS overnight at 4°C. Secondary antibodies were as follows: donkey anti-rabbit Alexa 488 (Abcam, ab150073); donkey anti-rat Alexa 594 (Abcam, ab150154); and donkey anti-guinea pig (Jackson Lab, 706475148). The next day, sections were rinsed three times for 10 min each in PBS, then mounted on SuperFrost glass slides (Thermo Fisher Scientific) in a petri dish with PBS, briefly rinsed with distilled water and coverslipped with mounting medium (Fluoromount-G, ThermoFisher). Finally, they were sealed with nail polish and stored at 4°C.

### Confocal Imaging

A LSM880 Zeiss Confocal Microscope was used to acquire tiled z-stack images of double hemisphere hippocampi using a Zeiss plan apochromat 20×/numerical aperture 0.8 objective. This was used to validate the injection and implantation site. Only the animals that had viral expression in both dorsal hippocampi were included in this study (see Figure 1A).

### Reinforcement learning (Q-learning)

Here, we sought to fit the behavior of mice performing the radial arm maze to reinforcement learning (RL) agents through a Q-learning approach [32]. For this comparison, we used the data from the first eight days (Trial block 1) of the behavioral experiments.

### Environment and Q-learning agent

In our setup, we modeled the maze as a discrete Markov decision process and defined ‘state’ as the current and previous location of the agent, as this modification enabled the agents to robustly perform entry sequences, a prominent feature of mouse behavior in the task (Figure 2A). There were 9 locations in total and the action selection was influenced by the values of the corresponding state-action pairs in the Q-table. These values were used in a softmax function to give the probability of choosing each action. The possible actions were dependent on the current state the agent occupied, save for a ‘stay’ action which was always possible. Beyond this action, the agent could either move to one of the arms from the center zone or return to this zone from one of the arms. The full state representation of the agent also included the previous state, so that the full state of an agent at time *t* is given by [*s_t−_*_1_*, s_t_*], where *s_t−_*_1_ is the previous state of the agent, and *s_t_* is the current state. Treat locations were assigned to the same three arms outlined in the experimental paradigm. The agent receives a reward *r*_treat_ = 1 when occupying these arms for the first time in each trial, and a negative reward *r*_else_ = −0.1 in all other cases. Furthermore, agents always began a trial in the bottom arm. Trials were terminated when all three treats were collected or when more than 500 actions were performed. Since the mice performed eight trials per day over eight days in Trial block 1 of the purely behavioral experiments, each agent performed 64 trials to match this quantity.

In the first trial, the Q-table was initialized with zeros in every element except for those whose stateaction pair corresponded to returning to an arm the agent occupied in the previous step. The latter was implemented to discourage immediate reversals. These Q-table values are used to determine the probability distribution for selecting each possible action in any given state. For example, for an agent in state [*s_t−_*_1_*, s_t_*] at time *t*, where the possible actions are contained in a set A, the probability of choosing some action *a_t_* ∈ A is given by

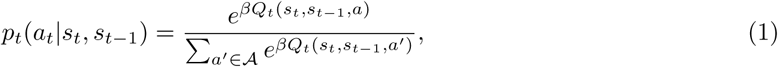

where *β* is inversely related to the degree of stochasticity in action selection. Supposing that action *a_t_* is selected, the Q-table is updated by the following:

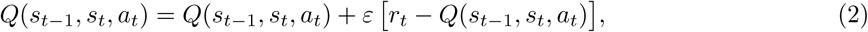

where *ε* controls the learning rate, and *r_t_* ∈ {*r*_treat_*, r*_else_}. Once the trial is completed, the final Q-table configuration is used as the initialization for the next trial, and this continues until all 64 trials are completed. For each trial, we calculate the number of errors that the agent made, which were entries into non-rewarded arms and re-entry into rewarded arms.

### Parameter estimation (fits of *ε* and *β*)

For each mouse, the optimal *ε* and *β* were estimated by minimizing the mean absolute error between model-predicted and observed daily mean errors across the first *D* = 8 days (Trial block 1 in the purely behavioral experiments), each comprising *T* = 8 trials. For a given parameter pair, we simulated *N* = 500 independent agents over the same *D* × *T* trials and computed the model-predicted daily mean error

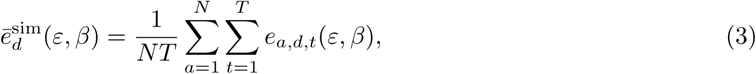

where *e_a,d,t_* is the error of agent *a* on trial *t* of day *d*. The parameters were then obtained as

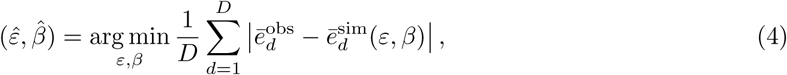

where 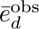 is the animal’s mean error on day *d*.

Optimization used SciPy’s differential evolution with strategy best2bin, tolerance of 0.1, population size 30, mutation constants (0.5, 1.0), recombination constant 0.7, Sobol initialization, deferred updates, and no polishing. Candidate solutions for *ɛ* and *β* were constrained between [0.001, 1] and [0.1, 20], respectively.

Once the optimal *ε* and *β* were estimated, we plotted the mean and standard error of the mean of the agent-averaged daily mean error using these parameters, overlaying the same for the corresponding mouse. Across mice, these *ε* and *β* values were plotted as scatter plots and individually as box plots separated by genotype with overlaid individual points. For the latter, significance was assessed using two-sided Mann–Whitney U tests (non-parametric, independent samples) and *p*-values were reported.

### Loss-surface and ‘area under the surface’ (AUS) analyses

Although the optimization provides an estimate of the optimal *ε* and *β* values for each animal, it does not directly inform us of how other values of epsilon and beta compare. Gaining this information can provide a general indication for the quality of the model itself in characterizing the behavior of the animals. Therefore, we also wanted to assess the quality of these fits over a range of parameters. This would give an indication for the appropriateness of the RL model in describing each mouse’s behavior in general. To do this, we computed per-mouse loss surfaces over a rectangular grid, where *ε* ∈ {0.1, 0.2, …, 1.0} and *β* ∈ {1, 2, …, 20}. For each grid point, we simulated 500 agents for 8 × 8 trials, converted trial-level errors to daily means, and computed the objective function against the mouse’s daily means. These loss values were plotted in 3D to visualize the loss surface. Further, this surface was summarized by its area under the loss surface (AUS), numerically integrating over the grid by Riemann sum with the known grid spacings. These AUS values were plotted as box plots separated by genotype and significance was assessed as described above.

### Noisy RL model variant

To supplement the AUS analysis, we also designed a noisy variant of the outlined RL model. In this model, we introduced additive zero-mean Gaussian noise to every element of the Q-table after every action. The standard deviation of this noise term is given by an additional hyperparameter, *q*_noise_. The higher the choice of *q*_noise_, the more unstable the learned behaviors become. To assess how this model compares to the original RL model, we repeated the AUS analysis, where *q*_noise_ was swept over the values (0.001–0.05 in 0.001 steps; 0.06, 0.07, 0.08, 0.09, 0.10, 0.15, 0.30), and AUS was calculated in each case for each mouse. These values were plotted per mouse and averaged over genotype, including the original RL model values also (which are equivalent to the noisy variant RL model with *q*_noise_ = 0). From this, we also determined the *q*_noise_ at which AUS was minimal for each mouse. These *q*_noise_ values were then displayed as box plots for each genotype and significance was assessed as described above.

### Data analysis and statistics

#### Behavioral metrics

Animals were tested in the RAM over 14 (behavioral experiments) or 6 (electrophysiological experiment) consecutive days. The maze and each arm were delineated into arms using either EthoVision XT (Noldus) or ImageJ (Fiji). To calculate the errors made by the animal per trial we considered the entry of each non-rewarded arm and the re-entry of a rewarded zone after reward consumption. Furthermore, if the animal entered a non-rewarded arm for the first time, it was considered a RM error, whereas re-entry into any arm was considered a WM error. Trails with zero errors are considered high-performance trials. Trial performance was averaged over days and compared across groups. Additionally, errors were analyzed in relation to reward locations by computing average errors made before obtaining the first, second, and third reward within each trial. This was also done for RM and WM errors.

To visualize trial-by-trial behavior, raster plots were generated showing the timing of entries into each arm. For each trial, the time of entry and the corresponding arm identity were recorded and color coded accordingly. Raster plots were aligned to trial start times, with each trial represented as a separate row. This representation allowed for a clear visualization of the sequence and distribution of arm visits across trials.

For each animal the trial speed was calculated from the recorded trajectories in the radial arm maze. Speed measures were derived for the entire trial and averaged over days. Additionally, speed was analyzed in relation to reward locations by computing average speed before the first, second, and third rewards within each trial. Trials were classified as high-performance (no errors) or error trials, and average speeds were compared between these conditions and across genotypes.

#### Rate maps

Spikes were speed-thresholded at 2.5 cm/s, and position was spatially binned into 1 cm × 1 cm bins within the 100 cm x 100 cm arena. Firing rates were computed by dividing the spike map by the corresponding occupancy map (time spent in each bin). Both maps were smoothed with a Gaussian kernel (*σ* = 2 cm) before normalization. Bins with no occupancy were excluded. Normalized rate maps were generated for each recording session (each day) and for each trial, respectively.

#### Spatial Information

The spatial information score, *SI*, was calculated for each cell based on the rate maps to measure how much information (in bits) each spike conveys about the animals spatial location. SI was calculated according to the Skaggs-McNaughton’s equation:

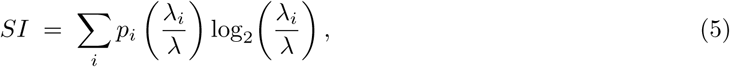

where *p_i_*is the probability of the animal occupying spatial bin *i* (occupancy fraction), *λ_i_* is the mean firing rate of the cell in bin *i*, and *λ* = Σ*_i_ p_i_λ_i_* is the overall mean firing rate across the entire environment.

#### Sparsity

The sparsity of each cell *S* ranges from 0 to 1. Values closer to 1 indicate that the firing is more evenly distributed across space, while lower values indicate that firing is concentrated to fewer spatial bins in a few locations (i.e. the cell is more spatially selective).

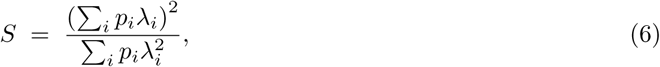

where *p_i_*is the probability of the animal occupying spatial bin *i* (occupancy fraction), and *λ_i_* is the mean firing rate of the cell in bin *i*.

#### Stability: Kullback-Leiber distance

To calculate the stability of this spatial firing, we adapted the equation for the Kullback-Leibler divergence. The KL distance (or relative entropy) is a metric used to quantify the difference in information represented by two distributions. In this case, the KL distance is calculated for each cell between trial *t* and trial *t* + 1. This gives a single cell measure for the change in spatial firing from one trial to the next. Suppose the firing rate in a spatial bin *i, j* in trial *t* is *λ_ij_* and in trial *t* + 1 it is *η_ij_*. Only spatial bins that were visited at least once in both *λ_ij_* and *η_ij_* were included for further analysis. If we consider a time bin of size Δ*t*, then the probability of getting *n* spikes in that spatial bin is:

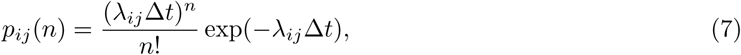

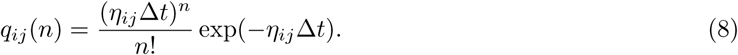

From these, we obtain

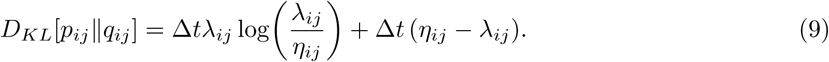

Since everything on the right-hand side is proportional to Δ*t*, we can just consider *D_KL_*[*p_ij_*||*q_ij_*]*/*Δ*t*. This is well-behaved, unless *λ_ij_ >* 0 but *η_ij_* = 0, that is, when the rate in a spatial bin (*i, j*) is zero. For smoothed rate maps, this only happens if the neuron in trial *t* + 1 has completely shut off. Such cases may be due to statistical fluctuations, since the neuron may have a very low but nonzero rate. To avoid this, we may use a regularization procedure, e.g. setting *η_ij_* to the minimum nonzero firing rate {*λ_kl_*} observed across all bins.

Using the above procedure, we can then define the change from trial *t* to *t* + 1 as

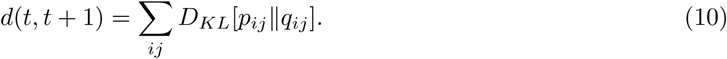

Note that the above is not symmetric, that is, *d*(*t, t* + 1) ̸= *d*(*t* + 1*, t*), because the KL divergence is not symmetric. We can, however, symmetrize this as

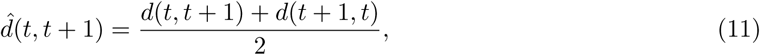

where *d*(*t* + 1*, t*) is defined similarly to *d*(*t, t* + 1) with *η* and *λ* replaced.

#### Reward raster plots and population vector analysis

To examine neuronal responses to reward delivery, spikes were aligned to the animal’s first entry into each of the three reward zones. Peri-event raster plots and histograms were constructed by aligning the spikes to reward entry and binning them in 20 ms time bins within a ±8 s window. Peri-event firing rates were calculated per bin, averaged across trials, and smoothed with a Gaussian kernel (*σ* = 5 bins). Neurons with mean firing rates exceeding 15 Hz were classified as putative interneurons and excluded from further analyses.

For each reward zone, peri-event time histograms were z-scored per neuron, and population activity matrices were generated. Neurons were sorted by the latency of their peak response, and this order was used to visualize heatmaps of z-scored activity around reward entry. To assess zone-specificity of the responses, neurons were sorted by their peak response to one zone.

Population vector (PV) analysis was then performed to quantify the similarity between reward zones. For each neuron, PV correlations were computed by correlating its peri-event firing rate vectors between all pairs of reward zones. At the population level, mean response vectors were calculated for each reward zone (zone × time bins), and Pearson’s correlation coefficients were used to generate zone-wise similarity matrices. PV correlations were computed separately for individual animals for each consecutive recording session (days 1–6).

#### Decoding

To assess whether neuronal ensembles contained information about reward and spatial location, we implemented two linear support vector machine (SVM) decoders using a one-vs.-rest scheme. For each session, spike trains from quality-controlled units were aligned to behavioral time, concatenated across trials, and binned at 40 ms resolution. Spike count matrices were trial aligned and labels were assigned as either reward vs. non-reward (binary) or reward zone identity (zones 1, 2, 3, and non-reward).

To improve spike coverage within trials, spike count vectors were smoothed with a Gaussian kernel (*σ* = 2 bins). Five-fold cross-validation was performed using trial blocks as folds. For each bootstrap iteration (100 replicates), a subsample of 17 neurons (which also served as a minimal inclusion threshold) was drawn with replacement and the decoding accuracy was computed relative to the true labels. Shuffle controls (1000 iterations per bootstrap) were used to establish chance performance.

Decoder performance was quantified as mean accuracy across bootstraps and trial blocks, with standard deviation as a measure of variability. Confusion matrices and receiver operating characteristic (ROC) curves were computed for both decoding modes to assess class-specific performance. ROC curves were used to visualize the trade-off between true-positive and false-positive rates for each class, providing a measure of the decoder sensitivity for each class label. Group-level analyses were performed by averaging decoding accuracy across animals and days, and comparing CR:DTA– and CR:DTA+ groups. Linear mixed-effects models were used to test for group × day interactions on decoding accuracy.

#### Statistics

Significance was measured as p<0.05 in all statistical tests. Group differences across sessions, days, or trials were primarily assessed with linear mixed-effects models, with group and time (day, session, or trial) and their interaction included as fixed effects and animal, subject, or neuron included as a random intercept; the significance of fixed effects was evaluated by ANOVA on the fitted model. Linear mixed-effects models were chosen because they account for repeated measurements within neurons across trials or days, allowing proper modeling of both within neuron and between-neuron variability while avoiding pseudo-replication.

Two-sided two-sample t-tests were used for simple pairwise group comparisons, and two-way repeated measures ANOVA (group × day) followed by post-hoc multiple comparisons was used for extended behavioral comparisons across testing phases. Non-parametric tests were used where appropriate: the Wilcoxon rank-sum test for between-group comparisons, the Friedman test for within-group comparisons across repeated conditions, and the Wilcoxon signed-rank test for post-hoc pairwise comparisons following a significant Friedman result. Binary pass/fail behavioral outcomes were additionally tested with Barnard’s exact test for 2×2 contingency tables, which was developed specifically for such categorical variables [60] adjusted for multiple comparisons using Nuisance parameter adjustment (Π). Population decoding was performed with a linear support vector machine, evaluated by 5-fold cross-validation and bootstrapped over 100 iterations with fixed cell sub-sampling. Chance-level performance was estimated via a label-shuffling permutation procedure, and classifier performance was further characterized using receiver operating characteristic (ROC) analysis.

## Acknowledgments

We thank B.A. Zaharia for technical assistance; H.M. Møllergård for genotyping; R.N. Raveendran and the Viral Vector Core Facility at the Kavli Institute for Systems Neuroscience, NTNU, for viral reagents; P.J.B. Girão for assistance with microscopy; and M.J. Nigro and members of the Quattrocolo laboratory for helpful discussions throughout the project. We also thank the animal facility staff at the Kavli Institute for Systems Neuroscience.

## Data availability

The code used for data analysis and figure generation in this study is publicly available on GitHub at https://github.com/IngvildGlaerum/CRc. Data generated in this study is available at: https://doi.org/10.18710/MURJBK

## Supplementary Information

**Figure Sup. 1:**
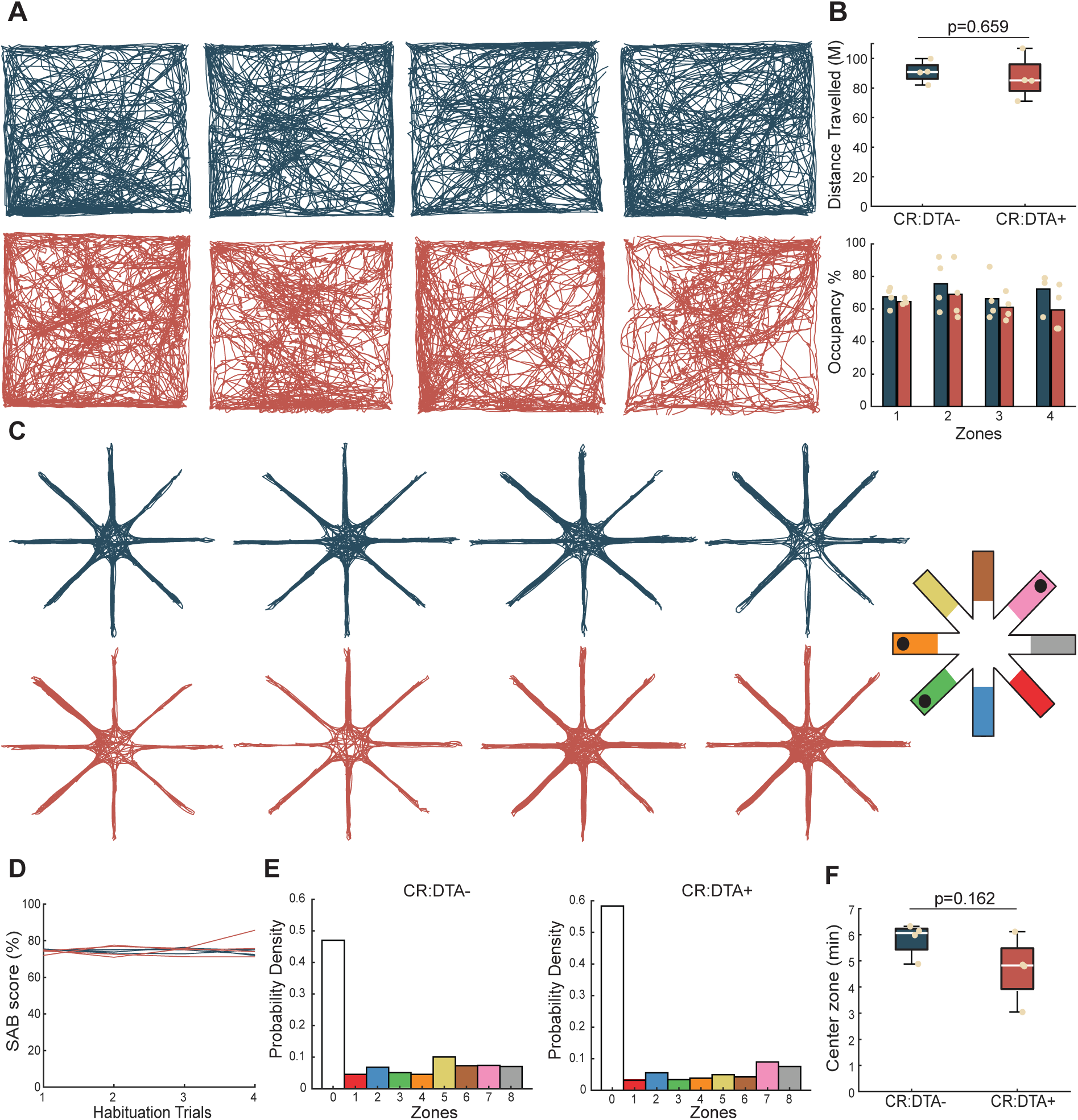
Exploratory behavior is not affected by CR cell ablation. **A:** Open field trajectory traces of all control animals (*top: blue*) and CR:DTA+ animals (*bottom: red*). **B:** No significant difference in the distance traveled during a open field exploration (*top*), with no difference in the occupancy of four equally spaced zones (*bottom*). **C:** Trajectory traces from RAM habituation trials for control animals (*top*) and CR:DTA+ animals (*bottom*). **D:** No significant difference in spontaneous alternation behavior (SAB) during the RAM habituation trials. **E**: No bias in the occupancy of any arm of the RAM, with the exception of the center in both controls (*left*) and CR:DTA+ animals (*right*). **F:** No significant difference in the time spent in the center zone between groups. Statistics: two-sample t-test

**Figure Sup. 2:**
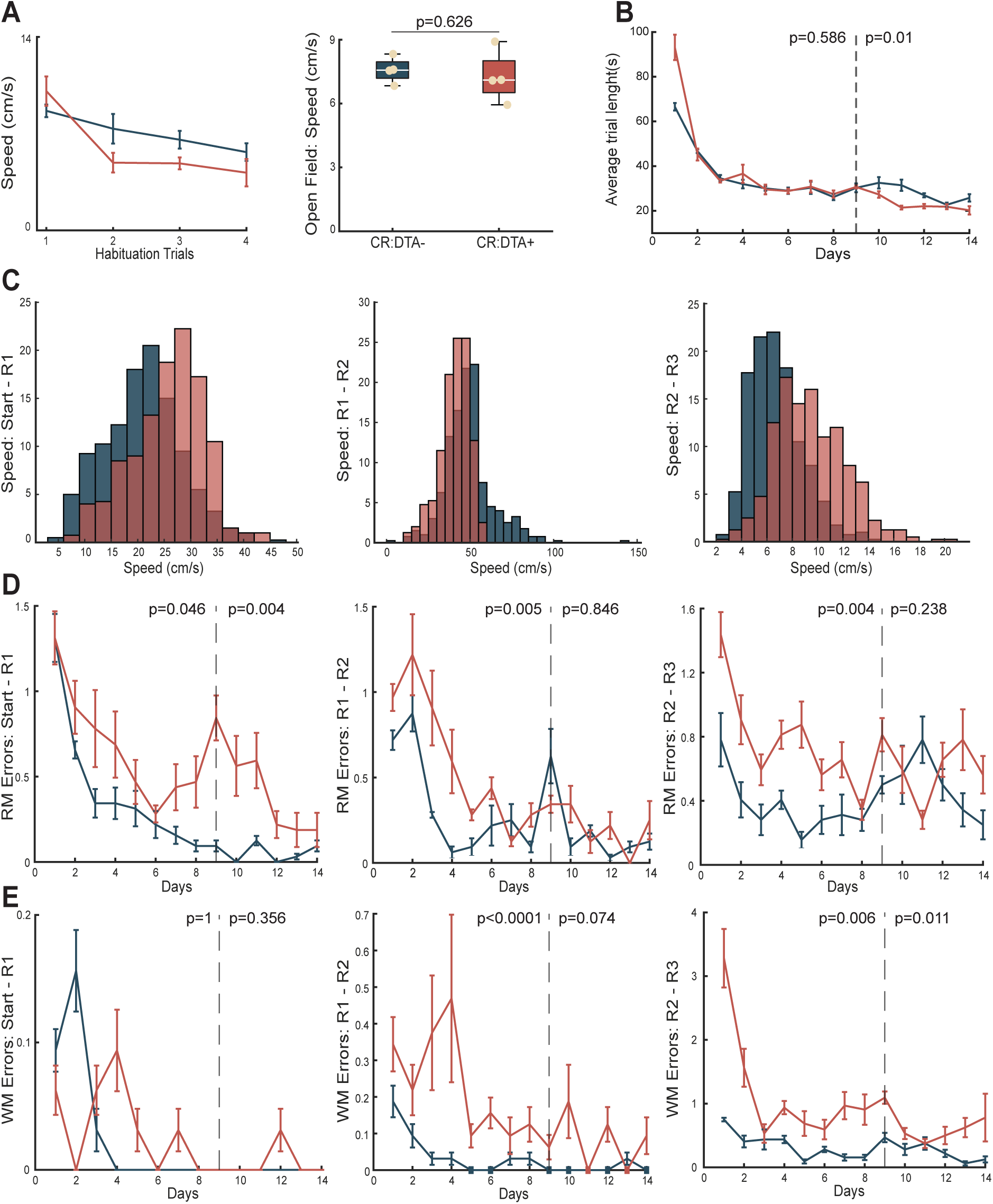
Speed and error types during goal-directed navigation. **A:** No significant difference in speed during habituation trials of the RAM (*left*), nor during open field exploration (*right*). **B:** No significant difference in trial duration in TB1, but controls had significantly longer trials in TB2. **C:** Speed distributions before obtaining R1 (*left*), R2 (*middle*), and R3 (*right*). **D:** Significant number of RM errors before obtaining R1 in TB2 (*left*), and a significant number of RM errors before R2 (*middle*) and R3 (*right*) in TB1. **E:** No significant number of WM errors before R1 (left), but a significant number of WM errors before R2 (*middle*) and R3 (*right*) in TB1. Statistics: repeated measures ANOVA or two-sample t-test.

**Figure Sup. 3:**
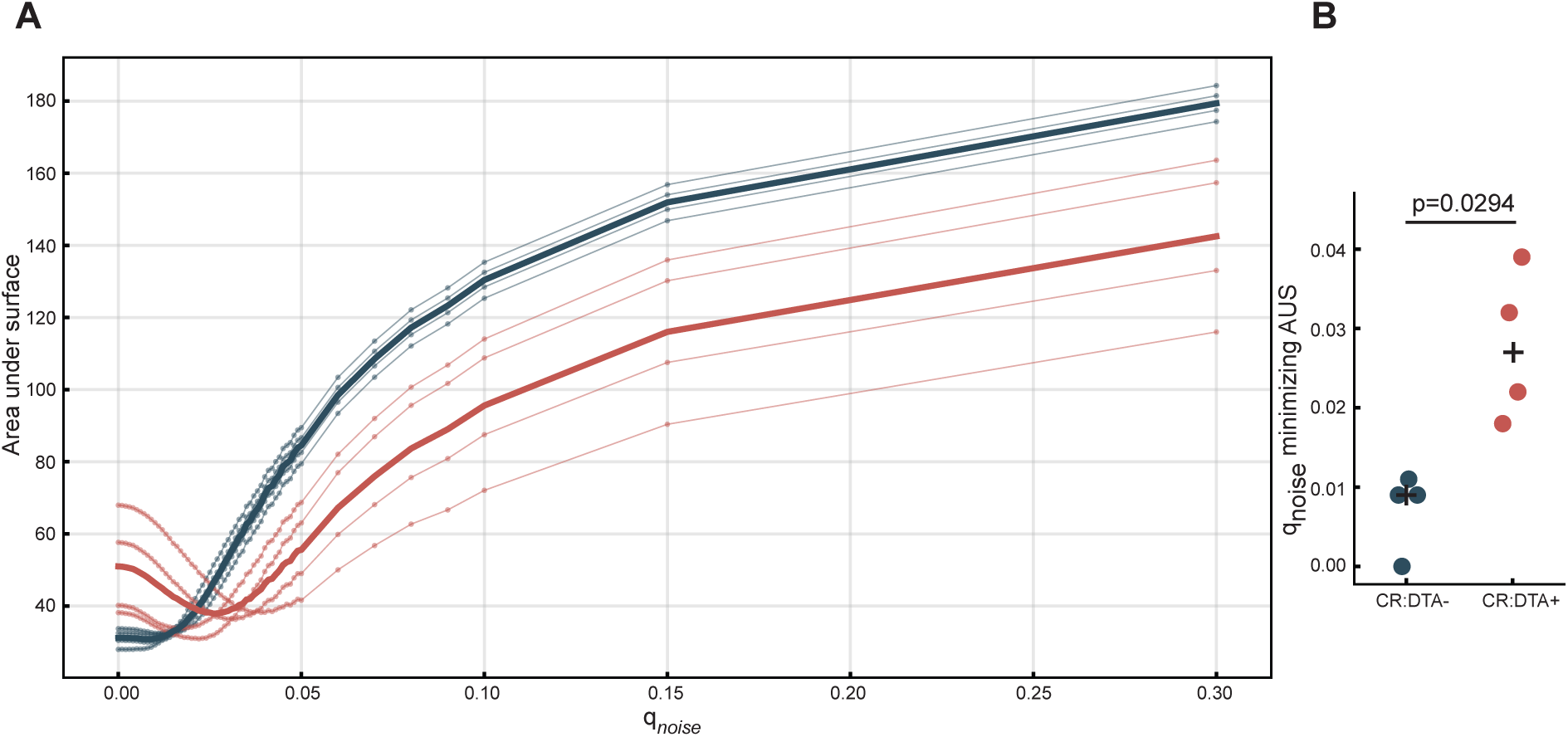
Introducing Gaussian noise to the Q-table **A**: Area under the loss surface for the noisy variant of the RL model over a range of *q*_noise_ values. **B**: Box plots for the value of *q*_noise_ that minimizes the area under the curve for each animal, separated by genotype. A plus symbol marks the average and significance were assessed using two-sided Mann–Whitney U tests.

**Figure Sup. 4:**
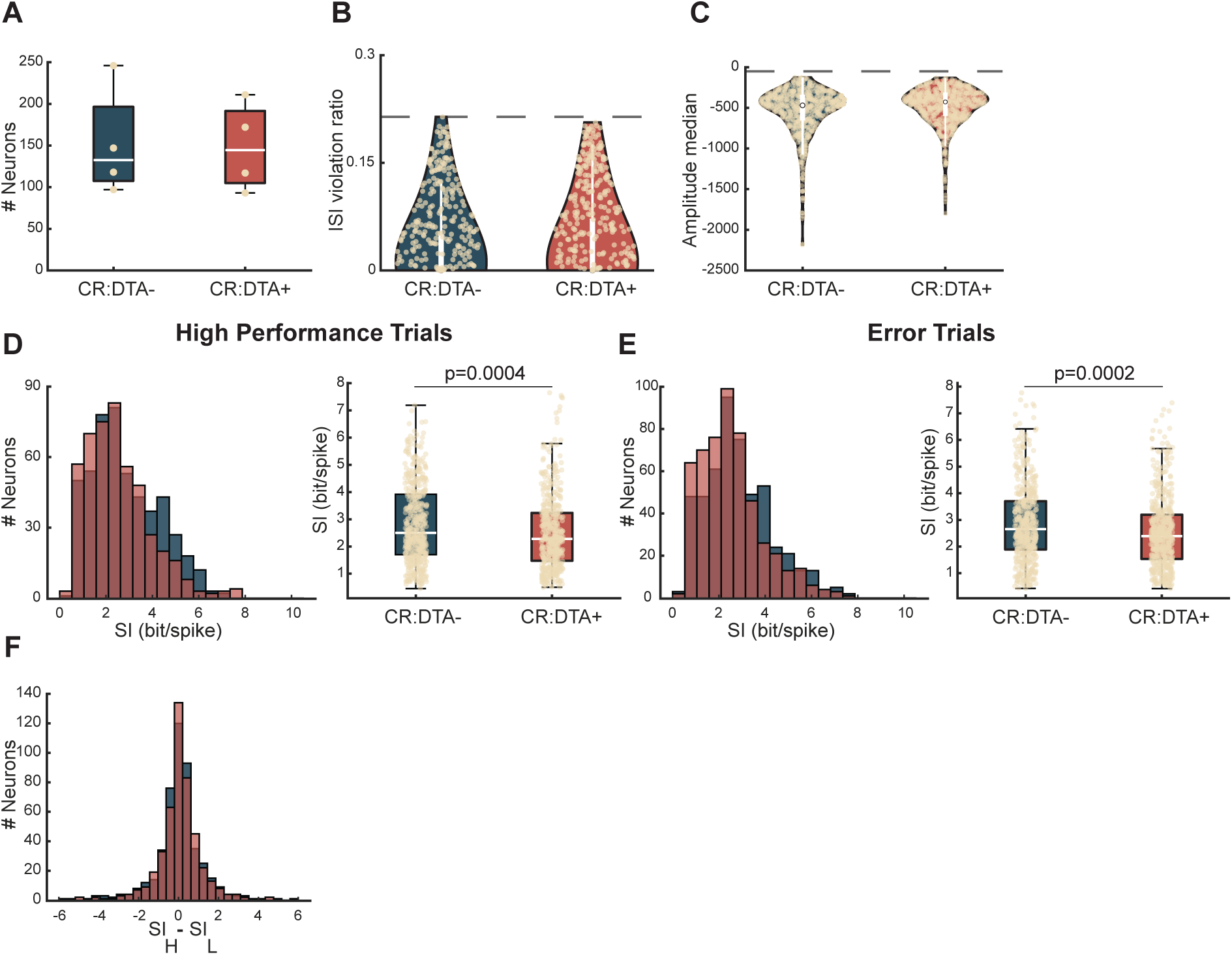
Additional single cell characteristics. **A**: Number of recorded single units per animal that passes the quality criteria in both groups. **B**: ISI-violation ratio per cell for included single units. Threshold at 0.2 (dotted line). **C**: Median Amplitude per cell for included single units. Threshold at -40*µ*V (dotted line). **D**: Spatial information (SI) calculated for exclusively high performance trials (no errors). Displayed as a distribution (*left*), and for single cells (*right*). **E**: Spatial information (SI) calculated for exclusively error trials (1 or more errors per trial). Displayed as a distribution (*left*), and for single cells (*right*). **F**: Ratio of SI for high performance trials (SI_H_) and error trials (SI_L_) calculated on a cell to cell basis. The individual cell does not change drastically in its spatial information content between high and low performance trials. Statistics: two-sided t-test or Wilcoxon rank-sum test.

**Figure Sup. 5:**
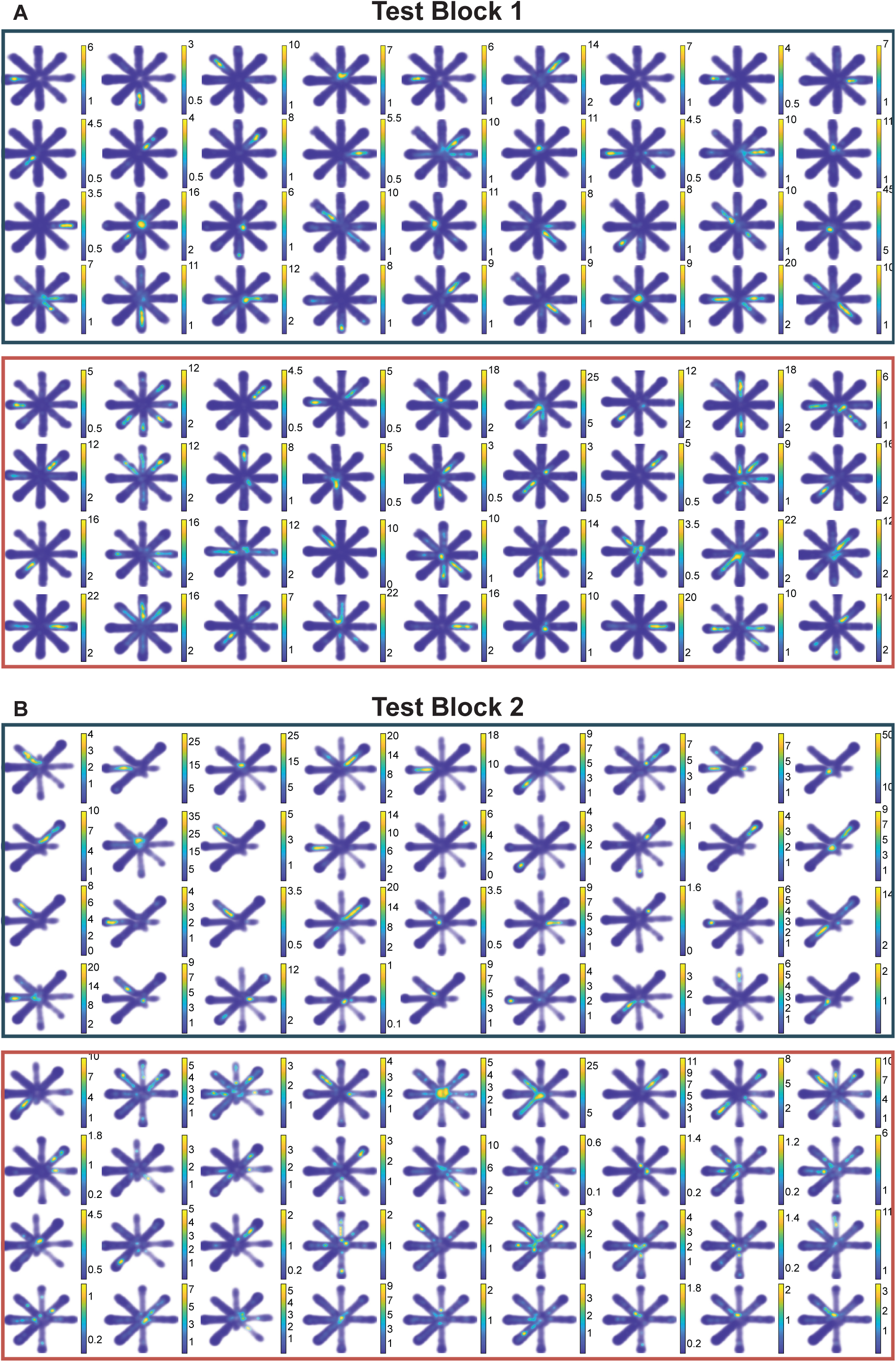
A larger sample of rate maps. **A**: Rate maps from Test Block 1 experimental day 1: CR:DTA– cells (*top*), and CR:DTA+ (*bottom*). **B**: Rate maps from Test Block 2 experimental day 4: CR:DTA– cells (*top*), and CR:DTA+ (*bottom*).

**Figure Sup. 6:**
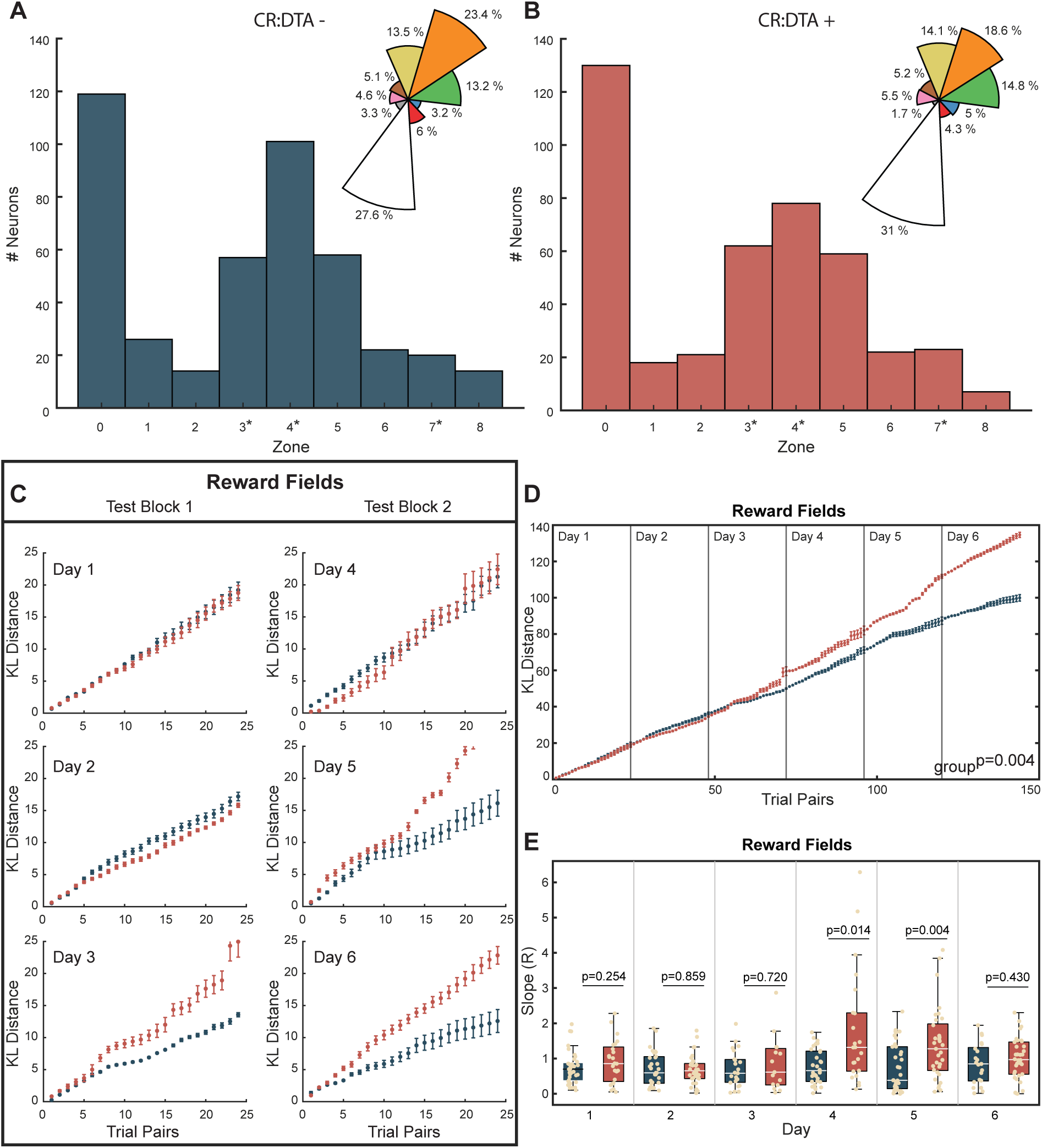
Neurons sorted according to where their field is located (coordinates of field peak). **A**: Distribution of field position according to zones for all cells in CR:DTA– animals. Nested figure displays a percentage representation of zones according to field location. **B**: Distribution of field position according to zones for all cells in CR:DTA+ animals. Nested figure displays a percentage representation of zones according to field location. 0 = center zone, asterisk indicating reward zones 1,2 & 3. **C**: Group averages of the KL distance calculated on executive trial pairs in each recording day in TB1 (*left*) and TB2 (*right*) for cells with a field in the reward zones. **D**: Continued cumulative median KL distance from each animal in calculated from cells with reward fields. **E**: Quantification of the slope (R: linear regression) of cumulative KL distance of each cell with a reward field for each day. Statistics: LMM (KL divergence ∼ group × session, animal as random effect).

**Figure Sup. 7:**
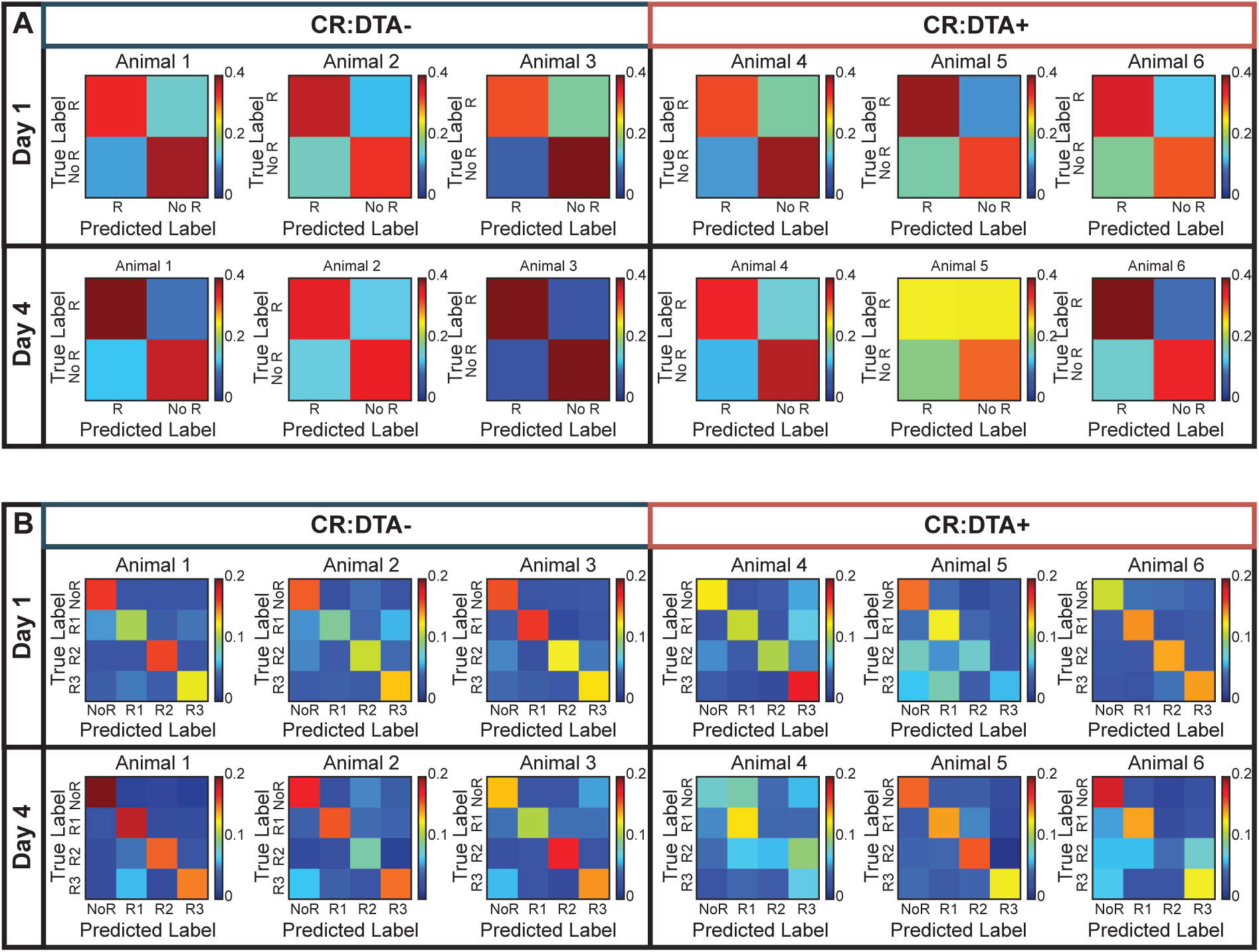
Decoding performance. **A**: Confusion matrices for decoding Reward vs Non-Reward for 3 example animals on days 1 and 4 for CR:DTA– (*left*) and CR:DTA+ (*right*). **B**: Confusion matrices for decoding Reward Zones for 3 example animals on days 1 and 4 for CR:DTA– (*left*) and CR:DTA+ (*right*).

